# SOX21 modulates SOX2-initiated differentiation of epithelial cells in the extrapulmonary airways

**DOI:** 10.1101/2020.06.04.134338

**Authors:** Evelien Eenjes, Marjon Buscop-van Kempen, Anne Boerema-de Munck, Lisette de Kreij-de Bruin, J. Marco Schnater, Dick Tibboel, Jennifer J.P. Collins, Robbert J. Rottier

## Abstract

SOX2 expression levels are crucial for the balance between maintenance and differentiation of airway progenitor cells during development and regeneration. Here, we describe SOX21 patterning of the proximal airway epithelium which coincides with high levels of SOX2. Airway progenitor cells in this SOX2+/SOX21+ zone show differentiation to basal cells, specifying cells for the extrapulmonary airways. We show that loss of SOX21 results in increased differentiation of progenitor cells during murine lung development. SOX21 inhibits SOX2-induced differentiation by antagonizing SOX2 binding on different promotors. SOX21 remains expressed in adult tracheal epithelium and submucosal glands, where SOX21 modulates SOX2-induced differentiation in a similar fashion. Using fetal lung organoids and adult bronchial epithelial cells, we show that SOX2+SOX21+ regionalization is conserved in human. Thus SOX21 modulates SOX2-initiated differentiation in extrapulmonary epithelial cells during development and regeneration after injury.

## INTRODUCTION

The lung is composed of a highly branched tubular system of airways lined with specific cell types, which together moisten, warm the air and filter out inhaled substances. The alveoli are located at the distal end of the airways. They consist of a thin layer of epithelium encircled by a network of blood vessels to facilitate an exchange of oxygen and carbon dioxide. During lung development, both growth and subsequent differentiation along the proximal-distal axis occur simultaneously. A well-regulated balance between progenitor cell maintenance, proliferation and differentiation is essential to ensure a fully functional lung at birth.

The formation of lung endoderm starts from the ventral anterior foregut by the development of two lung buds, forming the basis of the left and right lung. As the growing lung buds expand, the future trachea is separated from the esophagus proximal of the lung buds. Through a repetitive process of branching of the growing tips and outgrowth of the newly formed branches, a complex bronchial tree develops (Metzger, Klein et al. 2008) and regionalization of the branching structures occurs along the proximal-distal axis. Distal progenitors expressing the SRY-box protein SOX9 and the HLH protein ID2 generate new branches and will ultimately give rise to alveolar cells (Rawlins, Clark et al. 2009). While the buds grow and expand, SOX9+ progenitor cells gradually become more distant from the branch-inducing FGF10 signal secreted by the distal mesenchymal cells, resulting in loss of SOX9 expression and initiation of SOX2 expression (Park, Miranda et al. 1998, Weaver, Yingling et al. 1999). These SOX2+ progenitor cells mark the non-branching epithelium and give rise to the airway lineages.

SOX2 is a critical transcription factor in the development of the airways and epithelium lineages. Deficiency of SOX2 resulted in aberrant tracheobronchial epithelium due to the loss of basal cells (Rock, Onaitis et al. 2009, Tompkins, Besnard et al. 2009, Wang, Tian et al. 2013). In contrast, overexpression of SOX2 leads to an increase in basal cells and in addition to a defect in branching morphogenesis (Gontan, de Munck et al. 2008, Ochieng, Schilders et al. 2014). The requirement of proper SOX2 levels in airway development is well documented, but it is not clear how the balance between SOX2+ progenitor maintenance and early cell fate determination is regulated.

Previously, we showed that increased SOX2 expression during lung development leads to increased SOX21 expression in airway epithelium (Gontan, de Munck et al. 2008). While the function of SOX21 in lung development is not known, SOX21 was found to be enriched in SOX2+ stalk (differentiating bronchiole) versus tip progenitor cells of the human fetal lung (Nikolic, Caritg et al. 2017). Moreover, concomitant expression of SOX2 and SOX21 results in either a synergistic or antagonistic function depending on the tissue or environmental stimuli (Mallanna, Ormsbee et al. 2010, Freeman and Daudet 2012, Whittington, Cunningham et al. 2015, Goolam, Scialdone et al. 2016).

We hypothesized that SOX21 is an important regulator of SOX2+ progenitor maintenance during development and regeneration of airway epithelium. In this study, we demonstrate that SOX21 demarcates a region in the proximal-distal patterning of the airway tree during lung development. In this SOX21 positive region differentiation occurs of SOX2+ progenitor cells to basal cells and defines the epithelium of the extrapulmonary airways. Using heterozygous SOX2, SOX21 and full SOX21 knock-out mice, we show that variable levels of SOX21 antagonizes SOX2 function by suppressing the differentiation of progenitor cells during lung development. In the adult lung, co-expression of SOX2 and SOX21 is maintained in the extrapulmonary airways. Basal cells with reduced SOX21 levels loose stemness and are more prone to differentiate, similar to what is seen in proximal epithelial progenitor cells during lung development. Taken together, our results demonstrate that reduced levels of SOX2 or SOX21 compromise airway progenitor cell maintenance and differentiation during lung development and regeneration.

## RESULTS

### SOX2 and SOX21 regionalize the trachea and main bronchi in a proximal-distal pattern during branching morphogenesis

To gain more insight into the interplay between SOX2 and SOX21, we first examined the spatial and temporal distribution of SOX21 in lung epithelium with respect to the SOX2-SOX9 proximal-distal patterning at different stages of lung development.

The earliest expression of SOX21 during lung development was found at gestational age E11.5. A few cells expressing SOX21 were located in the most proximal region of the SOX2+ epithelium (Fig. 1A). This indicates that SOX2 precedes SOX21 expression which can be detected at E9.5 (Que, Okubo et al. 2007, Gontan, de Munck et al. 2008). At E11.5, abundant expression of SOX21 is seen in cells of the thymus and esophagus which also co-expressed SOX2. At E12.5, SOX21 expression became more apparent, but stayed restricted to the proximal part of SOX2+ epithelium, the trachea and main bronchi. From E13.5 onwards, SOX21 expression started to be expressed throughout the trachea and main bronchi, but was absent in the smaller SOX2+ airways. Four different zones in the developing lung epithelium could be distinguished throughout lung development (Fig. 1A): zone 1, the most proximal zone, the developing trachea and main bronchi, which consists of SOX2+ and SOX21+ airway epithelial cells; zone 2, which contains SOX2+ only airway epithelial cells; zone 3, a transition zone, in which distal SOX9+ progenitors transition into SOX2+ airway progenitors (Mahoney, Mori et al. 2014); and zone 4, the most distal part of the lung epithelium which contains the SOX9+ progenitors (Rawlins, Clark et al. 2009) (Fig. 1A). SOX21 was never observed in the distal buds and always in cells that also express SOX2. SOX21 is heterogeneously expressed between cells during the early developmental stages (E12.5-E14.5) and in the transition between zone 1 to zone 2 at later ages (>E15.5), in contrast to the homogeneous distribution of SOX2 (Fig. 1A). At E18.5, SOX21 remains expressed in the trachea, heterogeneously expressed in the main bronchi and absent in the intrapulmonary airways (Fig. 1A). Thus, SOX21 is expressed during the formation of the airway tree and is located in the extrapulmonary airways, which is the most proximal part of the SOX2+ epithelium.

**Figure 1.**
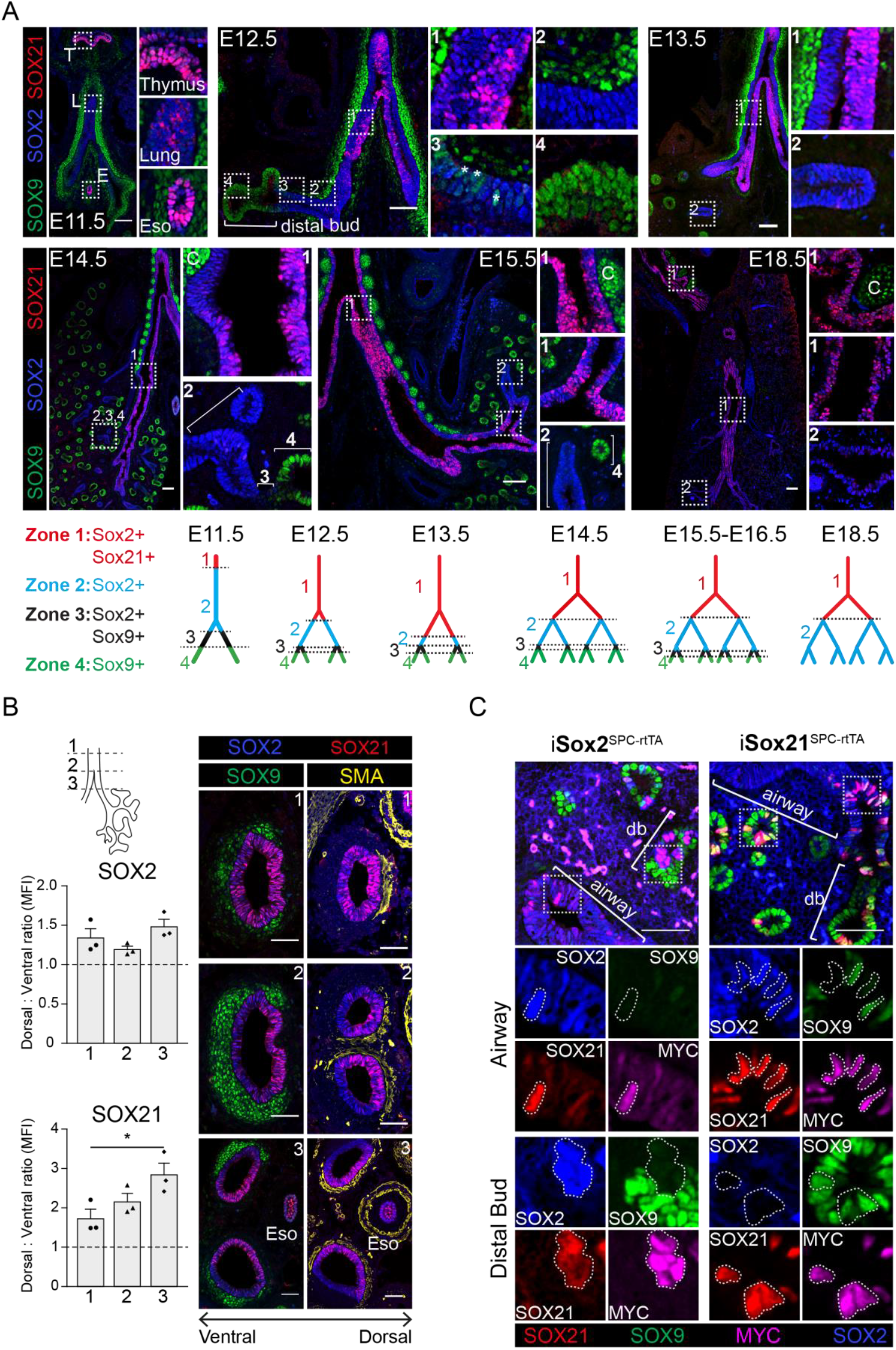
SOX21 is expressed in the airway epithelium and is confined to the proximal region of the SOX2 non-branching zone. (A) Co-staining of SOX9 (green) for distal buds, SOX2 (blue) for proximal epithelium and SOX21 (red) at different stages of lung development. Schematic representations show the distribution of SOX9, SOX2 and SOX21 in the different identified zones in the branching airways during gestation. At E12.5 the asterisks indicate the cells in zone 3 expressing both SOX2 and SOX9. T = thymus, L = Lung and E = Esophagus. C = Cartilage. Scale bar = 50 μm. (B) Transversal sections at three different locations of an E14.5 trachea and main bronchi stained with SOX2 (blue), SOX9 (green), SOX21 (red) and SMA (yellow). Location 1 = trachea, 2 = around the carina and 3 = immediately distal of the carina. SOX9+ mesenchymal cells surround the ventral side of the trachea and bronchi and SMA+ cells surround the dorsal side. The graphs show the ratio of mean fluorescence intensity (MFI) between ventral and dorsal of SOX2 and SOX21 at the three different locations. Scale bar = 50 μm. Eso = esophagus. (C) E15 Lung sections of doxycycline induced iSox2^SPC-rtTA^ and iSox21^SPC-rtTA^. Expression of transgenic *Myc*-tagged *Sox2* or *Sox21* was induced by giving doxycycline from E6 onwards. Sections are stained with SOX2 (blue), SOX9 (green), SOX21 (red) and MYC (purple). Scale bar = 50 μm. Db = distal bud.

In addition to this patterning, we observe a unilateral expression of SOX21 in the main bronchi, starting from the bifurcation of the trachea (Fig. 1A, S1A). SOX21 is almost absent at the lateral side, whereas SOX21 is abundantly expressed at the medial side of the airway (Fig S1A). Previous research showed that a higher expression of SOX2 was found in the dorsal tracheal epithelium compared to the ventral side of the tracheal epithelium during development (Que, Luo et al. 2009). In the developing trachea, SOX9+ cartilage nodules are found on the ventral side and Smooth Muscle Actin+ (SMA+) mesenchymal cells on the dorsal side (Hines, Jones et al. 2013). We made transverse sections from the trachea up to the main bronchi and observed a more abundant expression of SOX21 at the dorsal side, comparable to the location of high SOX2 levels (Fig. 1B). This unilateral expression pattern is present during branching morphogenesis but becomes less apparent after E15.5 (Fig. S1A). We hypothesized that the high expression of SOX2, specifically on the dorsal side of the trachea and main bronchi, is important for setting up the expression pattern of SOX21 during lung development.

To confirm whether increased levels of SOX2 induce expression of *Sox21*, we ectopically overexpressed a MYC-tagged SOX2 by using a tetracycline-inducible *Sox2* transgene under an SPC promoter-driven *rtTA* (Gontan et al, 2008) (Fig. S1B). SOX2 overexpressing cells showed a loss of SOX9 and an induction of SOX21 in the distal lung buds, thereby forcing distal progenitor cells to a proximal cell fate (Fig. 1C). Next, we generated a similar mouse model with a doxycycline inducible *Myc*-tagged *Sox21* transgene (Fig. S1B). Similar to the overexpression of SOX2, the induction of SOX21 results in the appearance of cystic structures, albeit much smaller (Fig. S1C). Further analysis showed that cells ectopically expressing SOX21 remained SOX9 positive and lacked SOX2 in the distal bud (Fig. 1C). Thus, SOX21 alone is not sufficient to induce a SOX2+ airway cell fate in the presence of distal mesenchymal signaling.

### SOX21 and SOX2 are co-expressed in a zone prone to progenitor cell differentiation

SOX21 marks a specific proximal region of the airway tree during development and we therefore asked the question whether this region is distinct from zone 2, which is only positive for SOX2. Early in development, at E11.5, SOX2 and SOX21 were both expressed in the lung, esophagus and thymus. At this embryonic age, the esophagus and thymus already contained TRP63+ epithelial cell progenitors (Fig. 2A), while TRP63+ basal cells only appeared in the lung at E12.5 (Fig. 2A, 2B). We previously showed that SOX2 directly regulates the expression of TRP63 (Ochieng, Schilders et al. 2014), and we therefore analyzed whether SOX21 plays a role in the differentiation of airway progenitor cells to basal cells. We found that basal cells appear from E12.5 onward in zone 1, but not in zone 2 (Fig. 2A, 2B). At E14.5, SOX2 and SOX21 are prominently expressed in epithelial cells in close proximity to SMA-expressing mesenchymal cells (Fig. 1). Consistent with previous findings (Que, Luo et al. 2009), we indeed found an increased percentage of basal cells at the medial compared to the lateral side and at the dorsal versus ventral side (Fig. 2B, S2A), suggesting that crosstalk between the mesenchyme and epithelium is necessary for combined SOX21 and SOX2 expression and subsequent basal cell differentiation (Fig. 2C). Additionally, in SOX2 or SOX21 overexpressing mice, zone 1 is extended. Within the extended zone, basal cells were present even in the absence of proximal mesenchymal-epithelial crosstalk, showing that induction of both SOX2 and SOX21 is sufficient for the differentiation of SOX2 progenitors to airway basal cells (Fig. 2D).

**Figure 2.**
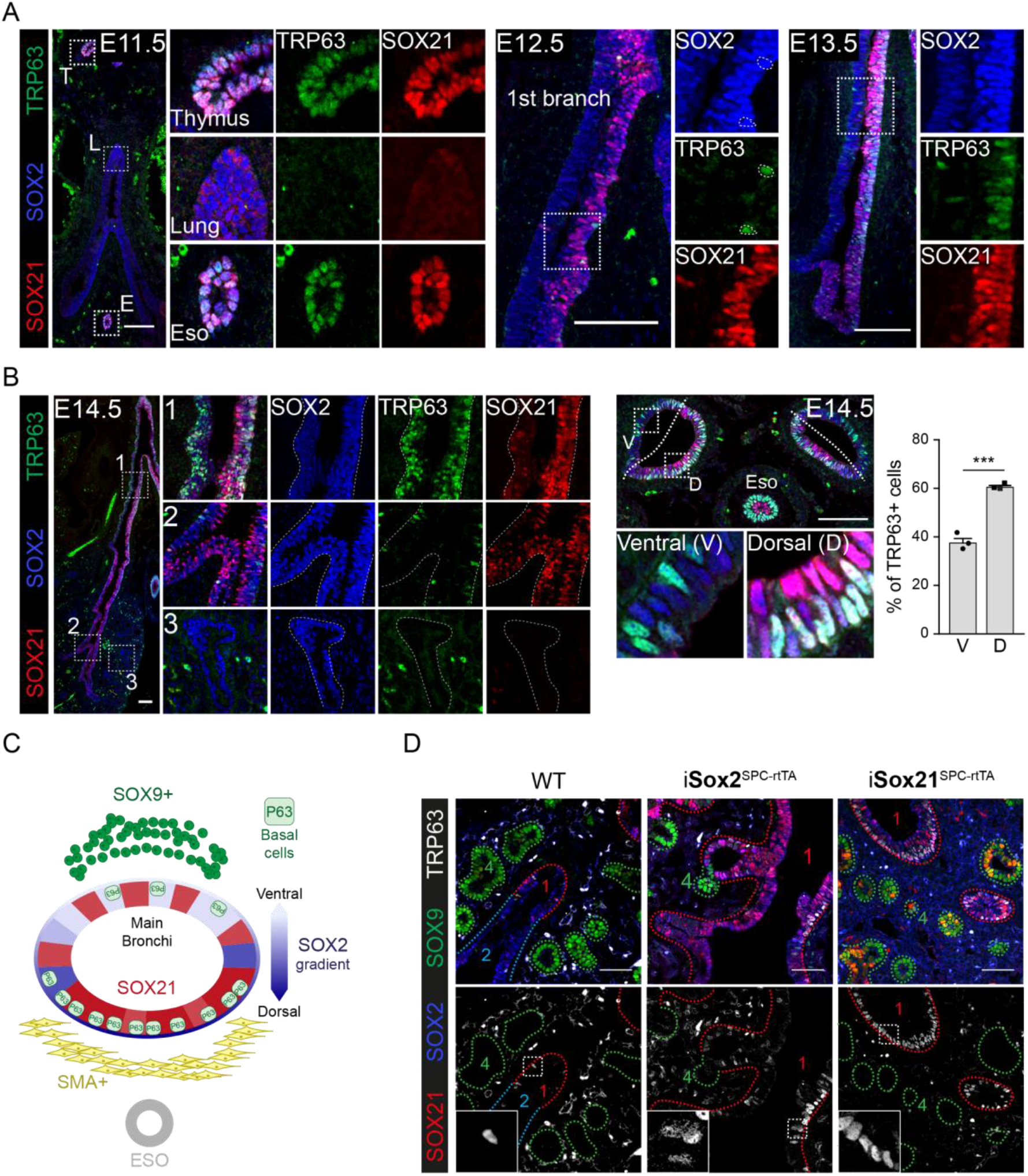
SOX2 and SOX21 are co-expressed in the zone where differentiation of progenitor cells to basal cells takes place. (A) Immunofluorescence of SOX2 (blue), SOX21 (red) and TRP63 (green) on lung sections of embryonic ages (E) 11.5, 12.5 and 13.5 showing a gradual co-localization of the three proteins. Scale bar = 100 μm. (B) Immunofluorescence of SOX2 (blue), SOX21 (red) and TRP63 (green) on longitudinal lung sections and transversal sections the main bronchi on embryonic ages (E) 14.5. The graph shows the percentage of basal cells in the ventral versus dorsal side of the bronchi. T-test (n=3; *p<0.05, *** p<0.001). V = ventral, and D = dorsal. Scale bar = 100 μm. (C) Schematic representation of SOX2, SOX21 expression and basal cell differentiation in a transverse section of a main bronchi. (D) E15 Lung sections of doxycycline induced (E6 until E15) iSox2^SPC-rtTA^ and iSox21^SPC-rtTA^ mice. Sections are stained with SOX2 (blue), SOX9 (green), SOX21 (red) and TRP63 (grey). Scale bar = 100 μm.

### SOX21 and SOX2 regulate maintenance and differentiation of the airway progenitor state

Next, we studied the effect of reduced levels of SOX21 on the differentiation to basal cells in *Sox21* heterozygous and homozygous knockout mice. *Sox21* knock-out (*Sox21*^-/-^) mice do not show respiratory distress at birth (Kiso, Tanaka et al. 2009). Their lungs are smaller than wild type littermate controls, but have no apparent branching defect (data not shown). However, an increased number of basal cells is present in *Sox21* heterozygous mice (*Sox21*^+/-^), which is even more pronounced in *Sox21*^-/-^ mice (Fig. 3A). Contrasting this increased number of basal cells, a decrease in the number of basal cells is observed in lungs of *Sox2* heterozygous mice (*Sox2*^+/-^) at E14.5 (Fig. S2B), corresponding with the dose-dependent role described for SOX2 in airway differentiation (Que, Okubo et al. 2007). Hence, SOX2 and SOX21 must have opposite effects. SOX21 maintains the SOX2 progenitor state and prevents differentiation, while SOX2 expression is important to initiate progenitor to basal cell differentiation.

**Figure 3.**
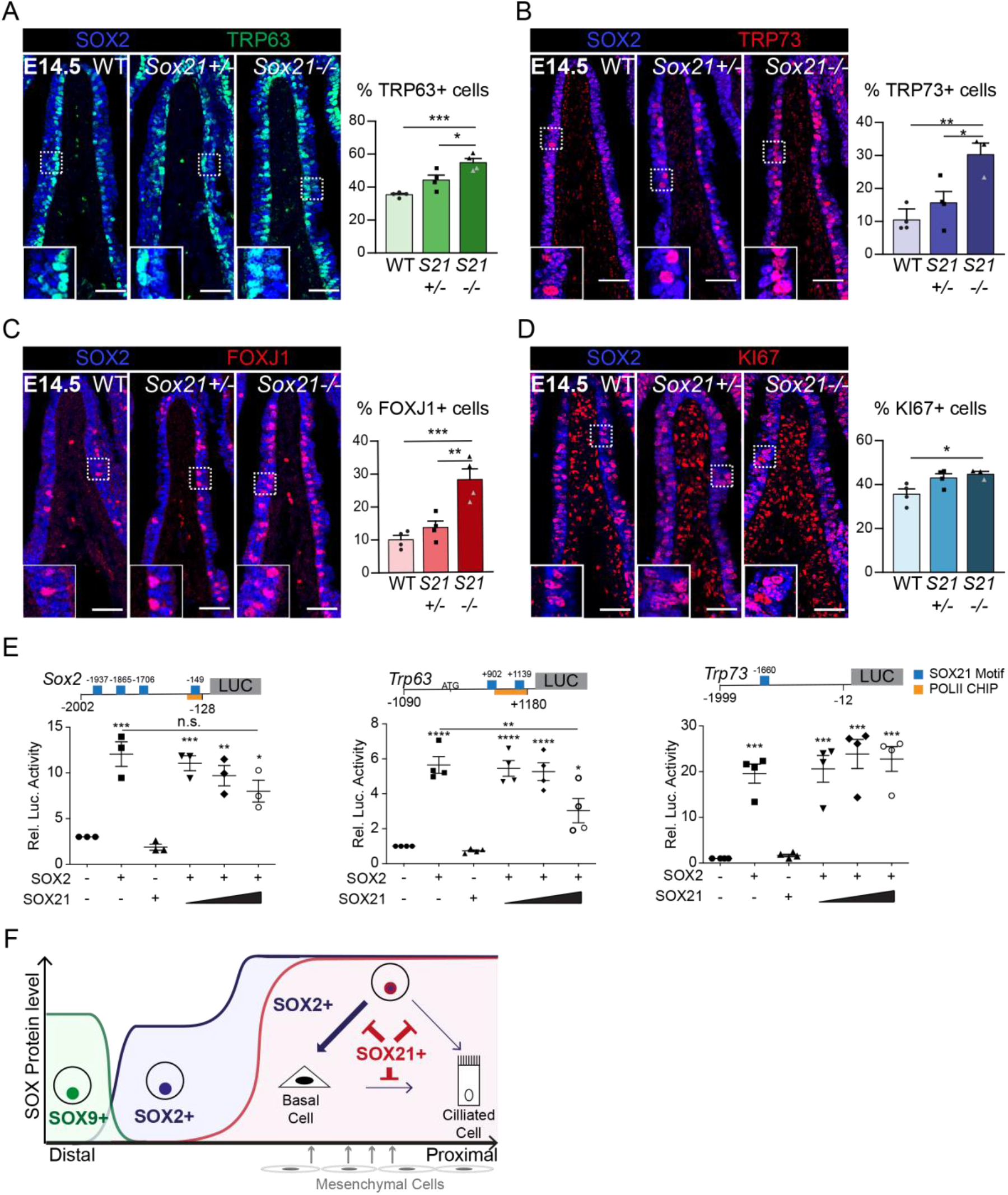
SOX21 counter balances SOX2+ progenitor differentiation to airway specific cell types. (A-D) Immunofluorescence and quantification of the number of TRP63+ basal cells (A), TRP73+ cells (B), FOXJ1+ ciliated cells (C) and KI67+ dividing cells (D) at E14.5 in wildtype (WT), *Sox21*+/- and *Sox21*-/- mice. The number of cells were counted within the first 400 μm immediately distal of the carina at the medial side of the airway. One-way ANOVA (n=3; * p<0.05). Scale bar = 50 μm. (E) Luciferase assay to test the transcriptional activity of a the *Sox2* promotor region from - 2002 till −128, *Trp63* promotor region from −1090 till +1180 and *Trp73* promotor region from - 1999 till −12 (+1 is considered the transcriptional start site). Blue squares showing SOX21 binding motifs and the orange bar shows the region bound by RNA polymerase II. The graph shows luciferase activity induced after transfection of FLAG-SOX2 and/or increasing amounts of MYC-SOX21 were co-expressed. One-way ANOVA (n=3; ** p<0.01, *** p<0.001, **** p<0.0001, when not indicated otherwise, stars show significance compared to control). (F) Schematic overview of epithelial progenitor maintenance and differentiation during murine lung development. From distal to distal a transition of SOX protein levels takes place. The most proximal part of the lung, the trachea and the two main bronchial branches, show high expression levels of SOX2, which initiates the expression of SOX21. SOX21 balances the maintenance of the SOX2 progenitor state in a zone where progenitor cells are prone to differentiate to airway specific cell types. High SOX2 levels stimulate differentiation.

To identify whether SOX21 and SOX2 levels are important for further differentiation to airway specific cell types, we also investigated the differentiation to ciliated cells. These start to appear at E14.5 and can be quantified using TRP73, an early marker for differentiation, and FOXJ1, a more mature marker (Marshall, Mays et al. 2016). At E14.5, we detect a small increase in the number of TRP73+ and FOXJ1+ ciliated cells present in the *Sox21*^+/-^ airways and an even larger increase in the *Sox21*^-/-^ airways (Fig. 3B, C). This again demonstrates that SOX21 levels are important for maintenance of the progenitor state and suppression of differentiation. In the *Sox2*^+/-^ mice, we observed no difference in the number of TRP73+ and FOXJ1+ ciliated cells (Fig. S2C, D), suggesting that decreased levels of SOX2 does not influence the initiation of ciliated cells despite that SOX2 can regulate the TRP73 promotor (Fig. 3E). Moreover, we did not observe changes in proliferation during development in *Sox2*^+/-^ airways and only a small increase in proliferation in the *Sox21*^-/-^ airways (Fig. 3D and Fig. S2E), showing that SOX2 and SOX21 act mainly on differentiation and not proliferation. Chromatin immunoprecipitation experiments have characterized SOX2 and SOX21 binding motifs and we identified putative binding sites for SOX21 in the promoter regions of *Sox2*, *Trp63* and *Trp73* (Matsuda, Kuwako et al. 2012) (Fig. 3E). To better understand the interaction of SOX2 and SOX21 during the initiation of differentiation towards ciliated cells, we performed luciferase assays, and show that SOX2 activates its own promoter, as well as the minimal promoter regions of the *Trp63* and *Trp73* genes (Fig. 3E, S2F). When increasing amounts of SOX21 were added to SOX2, we observed a significant decrease of luciferase activity with the *Trp63* promotor, a slight, but non-significant, decrease with the *Sox2* promotor and no difference with the *Trp73* promotor (Fig. 3E, S2F). This shows that SOX21 can suppress promotor regions of particular genes stimulated by SOX2. On the basis of these data, we propose that high levels of SOX2 initiate SOX21 expression, leading to a zone where basal cells start to differentiate. Within this zone, SOX21 promotes the maintenance of the SOX2+ progenitor state by inhibiting progenitors from differentiating, while SOX2 stimulates the differentiation of progenitors to basal cells (Fig. 3F).

### Deficiency of SOX2 decreases and SOX21 stimulates airway epithelial repair after naphthalene induced injury

Because the balance of SOX2 and SOX21 levels are important in maintaining a progenitor state during development, we investigated whether SOX21 is also important in the differentiation and maintenance of adult airway progenitor cells. SOX21 remains expressed throughout the tracheal epithelium (TE) in both basal and non-basal cells in adult mice (Fig. 4A). Basal cells are adult progenitor cells and are important for regeneration after injury (Rock, Onaitis et al. 2009). In addition, we observed SOX21 expressing cells in the submucosal glands (SMGs), which also regenerate the TE after injury (Lynch, Anderson et al. 2018, Tata, Kobayashi et al. 2018) (Fig. 4A). To test whether the levels of expression of SOX21 and SOX2 are important to control adult stem cell differentiation, we exposed wild type (WT), *Sox2*^+/-^ and *Sox21*^+/-^ mice to cornoil (CO) or naphthalene to induce transient epithelial injury. We examined the immediate response after 2 days and recovery after 5 and 20 days post-injury (DPI) (Fig. S3A, Fig. 4B). Due to fragility of the *Sox21*^-/-^ mice, we were unable to study adult *Sox21*^-/-^ TE after naphthalene injury (Kiso, Tanaka et al. 2009). Neither *Sox2*^+/-^ nor *Sox21*^+/-^ TE showed significant differences in the number of basal, dividing basal and ciliated cells when compared to WT after cornoil exposure (Fig. S3D, E). This suggests that the observed decreased and increased number of basal cells at E14.5 in *Sox2*^+/-^ and *Sox21*^+/-^ airways respectively (Fig. 3), is a transient delay in differentiation that is resolved after maturation of the lung.

**Figure 4.**
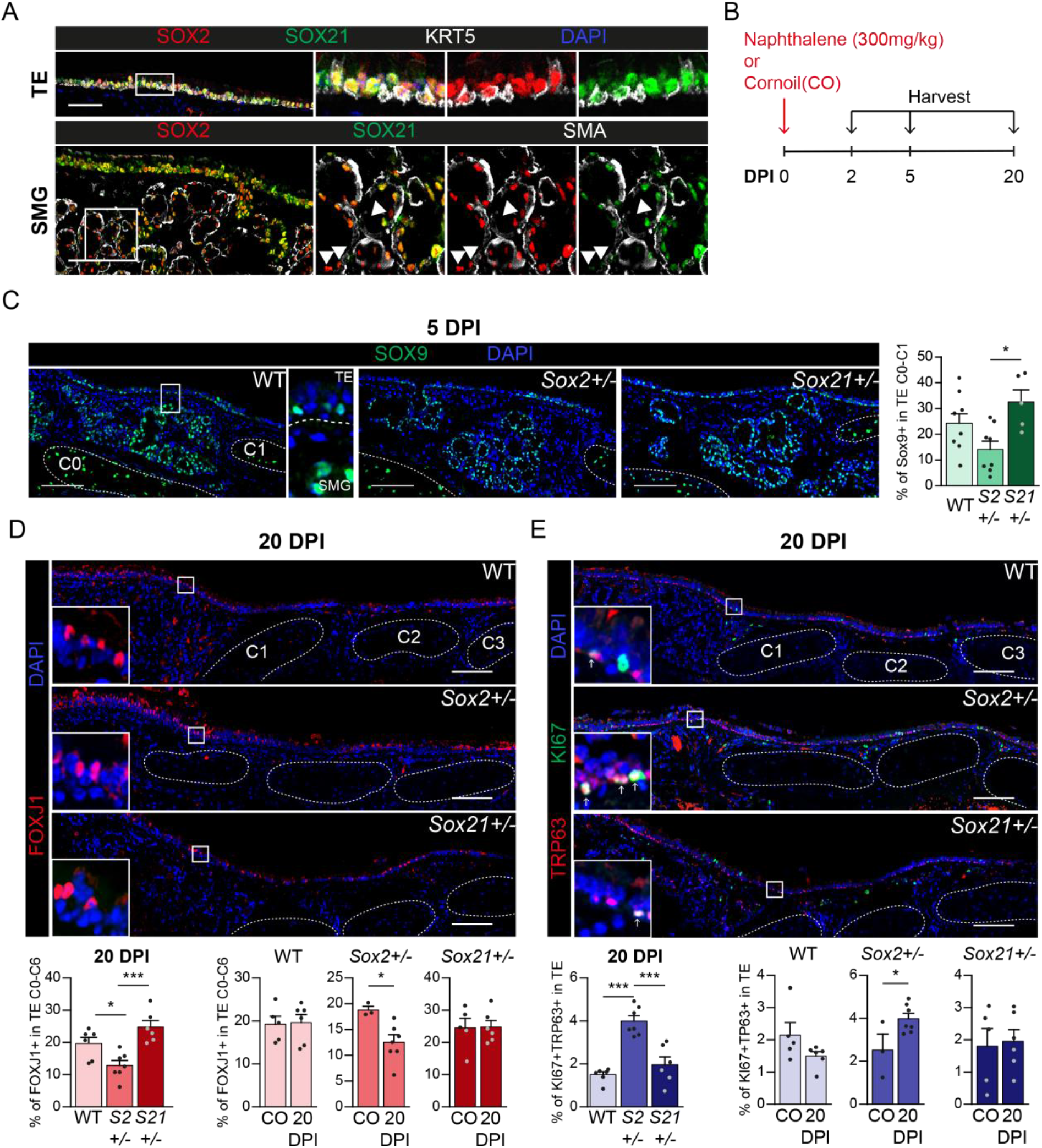
Regeneration is delayed in SOX2 deficient tracheal epithelium. (A) Immunofluorescence staining on tracheal sections of adult wild-type mice for SOX2 (red), SOX21 (green) and KRT5 (grey, top row) or smooth muscle actin (SMA, bottom row). TE = tracheal epithelium. SMG = submucosal gland. Closed arrowheads (►) indicate single SOX2+ cells. Scale bar = 100 μm. (B) Schematic overview of the experimental set up of the Naphthalene injury and recovery in wildtype (WT), *Sox2*+/- (*S2*+/-) and *Sox21*+/- (*S21*+/-) mice. (C) Immunofluorescence and quantification of the number of SOX9+ cells, 5 days post injury (DPI), in the upper TE from Cartilage (C) ring 0 till C1 in WT, *Sox2+/-* and *Sox21*+/- mice. Scale bar = 100 μm. One-way ANOVA (WT n = 8, *Sox2*+/- n = 8, *Sox21*+/- n = 5, * p<0.05) (D) Immunofluorescence of FOXJ+ (red) cells in the TE, of WT, *Sox2+/-* and *Sox21*+/- mice at 20 DPI. Scale bar = 100 μm. Quantification of the number of FOXJ1+ ciliated cells from C0 till C6 between genotypes. One-way ANOVA (WT n = 6, *Sox2*+/- n = 7, *Sox21*+/- n = 6, * p<0.05, *** p<0.001). Quantification of the number of ciliated cells, 20 DPI compared to the cornoil (CO) exposed mice of each WT, *Sox2*+/- and *Sox21*+/- mice. T-test (WT: CO n = 5 and 20DPI n = 6, *Sox2*+/-: CO n = 3 and 20DPI n = 7, *Sox21*+/-: CO n = 4 and 20DPI n = 6, * p<0.05). (E) Immunofluorescence with TRP63 (red) and KI67 (green) to mark dividing basal cells in the TE, 20 DPI, in WT, *Sox2*+/- and *Sox21*+/- mice. Scale bar = 100 μm. Quantification of the number of dividing basal cells from C0 till C6 between genotypes. One-way ANOVA (WT n = 6, *Sox2*+/- n = 7, *Sox21*+/- n = 6, *** p<0.001). Quantification of dividing basal cells, 20 DPI compared to CO mice of each wildtype (WT), *Sox2*+/- and *Sox21*+/-. T-test (WT: CO n = 5 and 20DPI n = 6, *Sox2*+/-: CO n = 3 and 20DPI n = 7, *Sox21*+/-: CO n = 4 and 20DPI n = 6, * p<0.05).

SOX9+ SMG cells surface at the TE after administering a high dose of naphthalene and thereby contribute to the repair of the TE. It was shown that at 1 DPI, SOX2 expression is largely extinguished from the SOX9+ SMG cells, suggesting that changes in expression are important in the early contribution of SMG contribution to repair (Lynch, Anderson et al. 2018). We were unable to detect differences in SOX2 or SOX21 expression at 2 DPI compared to cornoil exposed mice, suggesting that down-regulation of SOX2 at 1 DPI is temporary and a consequence of naphthalene administration (Fig. S3B). After five days, we observed a decrease of SOX9+ cells in *Sox2*^+/-^ TE, and a small increase of SOX9+ cells in *Sox21*^+/-^ TE compared to WT and to each other (Fig. 4C). This shows that the protein levels of both SOX2 and SOX21 are important for the regeneration of TE via the SOX9+ SMG cells. The number of ciliated cells and dividing basal cells was unaltered at 5 DPI (Fig. S3C).

To determine whether SOX2 or SOX21 deficiency affects regeneration after naphthalene injury, we quantified the percentage of ciliated, non-dividing and dividing basal cells at 20 DPI. Fewer ciliated cells were observed in *Sox2*^+/-^ TE compared to wild type and *Sox21*^+/-^ TE (Fig. 4D, S3D). Also, *Sox2*^+/-^ TE contained more dividing and non-dividing basal cells when compared to WT TE, or to *Sox2*^+/-^ cornoil exposed mice (Fig. 4E, S3D, F). In both WT and *Sox21*^+/-^ TE there were a similar number of ciliated cells, non-dividing basal cells and dividing basal cells at 20 DPI compared to cornoil exposed mice (Fig. 4D, Fig. S3D). These results indicate a recovery of injury in the WT and *Sox21*^+/-^ TE, while *Sox2*^+/-^ TE show a delayed recovery (Fig. 4E, S3D, F).

### SOX2 drives and SOX21 represses basal cell differentiation to ciliated cells

*Sox21*^+/-^ mice did not show increased basal or ciliated cell differentiation 20 days after naphthalene induced injury, contrary to the phenotype of increased differentiation we observed during development. To better understand the relation between SOX2 and SOX21 in the regulation of differentiation of basal cells an *in vitro* differentiation air-liquid interface (ALI) culture method of murine tracheal epithelial cells (MTECs) was applied. ALI-cultures provide standardized conditions to study basal cell differentiation at several different time points. Using this model, we show that SOX2 and SOX21 were both expressed at baseline levels at the start of ALI and their levels gradually increased in the initial days of ALI, something we were unable to demonstrate *in vivo* (Fig. 5A). Genes corresponding to the differentiation of ciliated and secretory cells are increased in a similar fashion (Fig. S4A). Immunofluorescence analysis of SOX2 and SOX21 on ALI day 10 showed expression of SOX2 and SOX21 in basal and luminal cells, but both proteins seem to have a heterogeneous distribution between cells (Fig. 5B). To determine whether the levels of SOX2 and SOX21 correlated with each other, we measured the fluorescence intensity of both proteins (Fig. S4B). Cells expressing SOX2 highly were mostly high in expression of SOX21 and vice versa (Fig. S4B). Basal cells were mainly Sox2^low^Sox21^low^, while ciliated cells were either SOX2^high^SOX21^high^ or SOX2^high^SOX21^low^ (Fig. 5C). Within TE *in vivo*, levels of SOX2 and SOX21 are similar between cell types (Fig. 4A), suggesting that SOX2 and SOX21 levels are stable for the maintenance of TE epithelium, but when differentiation is initiated both increase (Fig. 5G).

**Figure 5.**
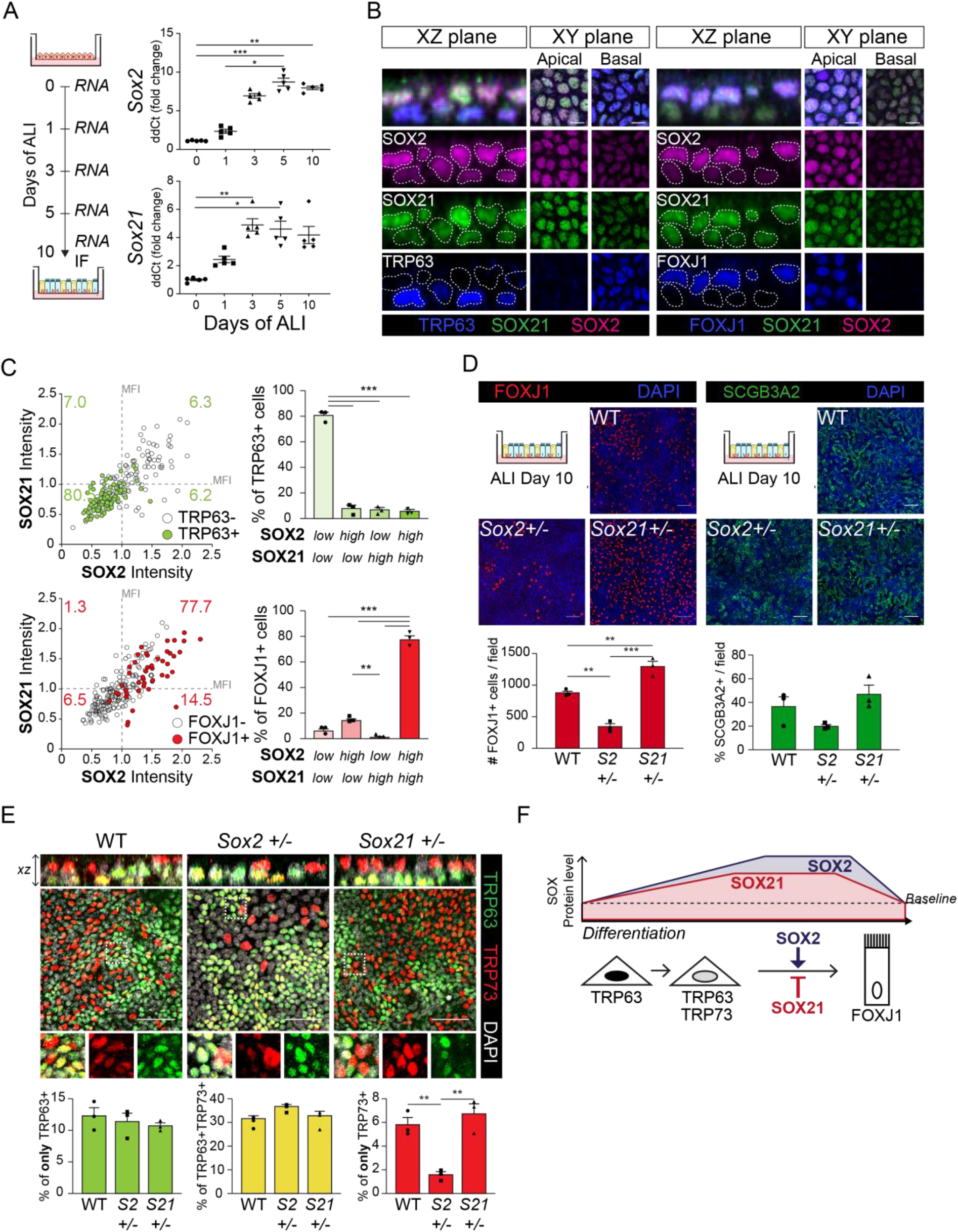
SOX2 and SOX21 are inversely correlated with basal cell differentiation to ciliated cells. (A) Schematic overview of mouse tracheal epithelial cell (MTEC) culture. QPCR analysis of *Sox2* and *Sox21* expression during differentiation of MTECs on Air-Liquid Interface (ALI). One-way ANOVA (n=5; * p<0.05, ** p<0.01, *** p<0.001). (B) Immunofluorescence staining on MTEC 10 days after ALI of SOX2 (purple), SOX21 (green), and TRP63 (blue, left images) or FOXJ1 (blue, right images). Dotted lines show representation of measured nuclei for fluorescence intensity. Scale bar = 25 μm. (C) Dot-plot of the MFI of SOX2 (x-as) and SOX21 (y-as). The green filled circles are TRP63+ basal cells and the number in each quadrant represents the percentage of basal cells in each quadrant. Bar graph is the quantification of basal cells that are either high (MFI>1) or low (MFI<1) in expression of SOX2 and SOX21. The red filled circles are FOXJ1+ ciliated cells and the number in each square represents the percentage of ciliated cells in each quadrant. Bar graph is the quantification of ciliated cells that are either high (MFI>1) or low (MFI<1) in expression of SOX2 and SOX21. One-way ANOVA (n=3; ** p<0.01, *** p<0.001). (D) Analysis of MTEC cultures of tracheal cells derived from wildtype (WT), *Sox2*+/- or *Sox21*+/- animals after 10 days of ALI culture. Immunofluorescence staining of ciliated cells (FOXJ1, red; C) or secretory cells (SCGB3A2, green; D). Scale bar = 50 μm. Quantification of the number of ciliated cells per 775×775 μm field. One-way ANOVA (n=3; ** p<0.01). (E) Analysis of MTEC cultures of tracheal epithelial cells derived from wildtype (WT), *Sox2*+/- or *Sox21*+/- animals, after 10 days of ALI culture. Immunofluorescence staining of TRP63 (green) and TRP73 (red). Scale bar = 50 μm. Quantification of the percentage of TRP63+ only basal cells, TRP73+ only cells, and TRP63+TRP73+ double positive basal cells. One-way ANOVA (n=3; ** p<0.01). (F) Schematic overview of in vitro basal cell differentiation. SOX2 and SOX21 are expressed at basal levels but upon differentiation stimuli, expression levels increase. Both proteins are highest expressed in luminal cells and ciliated cells. We propose a model, where SOX2 stimulates the transition of basal to ciliated cells, while SOX21 acts as an inhibitor in this process. Due, to the presence of FOXJ1+ SOX2^high^SOX21^low^ cells we propose a sooner downregulation of SOX21 to baseline levels.

To test the importance of the levels of SOX2 and SOX21 in basal cell differentiation, we isolated MTECs from WT, *Sox2*^+/-^ and *Sox21*^+/-^ mice. We observed decreased differentiation to ciliated cells in the *Sox2*^+/-^ and increased differentiation from basal to ciliated cells in *Sox21*^+/-^ (Fig. 4D). The differentiation capacity towards secretory cells was not affected in either *Sox2*^+/-^ and *Sox21*^+/-^ MTECs when compared to WT (Fig. 4D). Thus, the levels of SOX2 and SOX21 are mainly involved in balancing the differentiation to ciliated cells, with SOX2 stimulating differentiation and SOX21 inhibiting differentiation.

As we observed an increase in TRP73+ cells during airway development in SOX21^-/-^ mice, we assessed TRP73 expression in our MTEC cultures. TRP73 is one of the earliest markers expressed by basal cells upon initiation of ciliated cell differentiation, followed by the loss of TRP63 expression (Marshall, Mays et al. 2016). After 10 days of ALI there were comparable numbers of basal cells in *Sox2*^+/-^, *Sox21*^+/-^ and WT MTECs (Fig. S4E). While there was an increase in FOXJ1+ ciliated cells in *Sox21*^+/-^, neither the TRP73+/TRP63+ or TRP73+/TRP63-populations were affected (Fig. 5E, Fig. S4C). Furthermore, the number of TRP73+/TRP63+ population was also not affected by decreased levels of SOX2 (Fig. 5E, Fig. S4C). We did observe a significant decrease of single TRP73 expressing cells in *Sox2*^+/-^ MTECs. As both increasing levels of SOX2 and SOX21 are important to respectively boost and inhibit differentiation, it seems that both SOX2 and SOX21 act on ciliated cell maturation, but not on the initial induction of basal cell differentiation by the induction of TRP73 (Fig. 5F).

### SOX2 and SOX21 Expression is Conserved during Human Airway Epithelial Cell Differentiation

So far, our data show that SOX21 is an important factor for maintaining a balance between progenitor maintenance and differentiation in mouse lung development and regeneration. Next, we asked the question whether this is evolutionary conserved. Therefore, we cultured human fetal lung tip organoids as described previously (Nikolic, Caritg et al. 2017). While all cells were positive for SOX2, they co-expressed either SOX9, SOX21 or both (Fig. 6A). We observed that some of the cells expressing SOX21 were also positive for the TP63 basal cell marker in the fetal lung organoids (Fig S5A). To see whether lung progenitor cells expressing SOX21 are more likely to develop into a specific airway cell type, we differentiated the fetal lung organoids to airway organoids (Fig. S5A). After differentiation, the organoids contained both ciliated and basal cells and SOX21 expression was observed in most cells (Fig. S5A).

**Figure 6.**
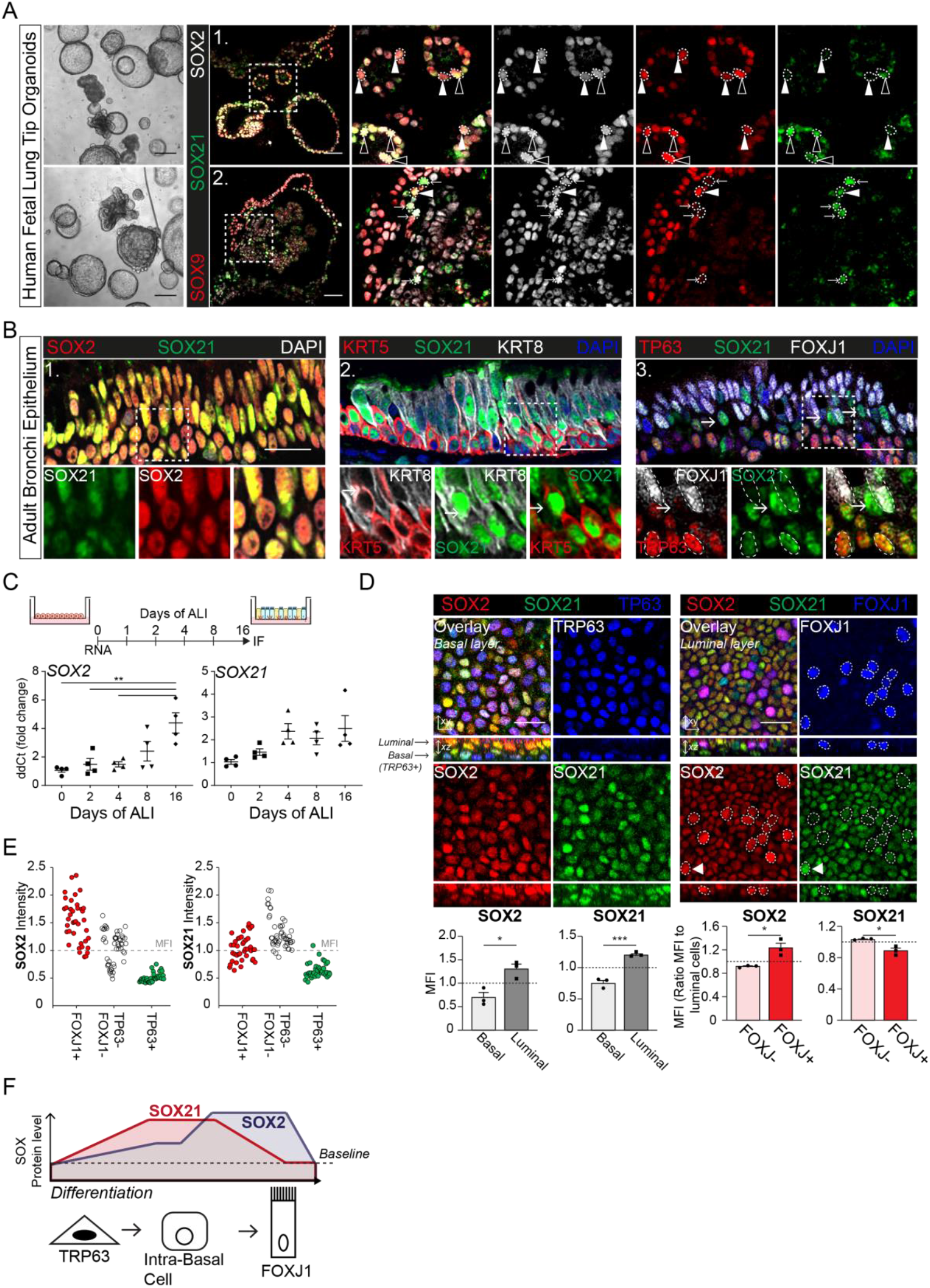
Evolutionary conserved expression of SOX21 in human airway epithelium. (A) Bright field images of human fetal lung tip organoids (scale bar = 250 μm) and immunofluorescence analysis shows SOX2 (grey), SOX9 (red) and SOX21 (green) positive cells in fetal lung organoids. Closed triangles (►) show cells positive for SOX2 and SOX9, open triangles (►) show cells positive for SOX2, SOX21 and SOX9. Arrows (→) show cells positive for SOX21 and SOX2. Scale bar = 50 μm. (B) Immunofluorescence analysis of sections of human adult bronchi shows co-localization of SOX2 (red) and SOX21 (green) throughout the epithelium (1). SOX21 (green) is expressed in luminal (KRT8; grey) and basal (KRT5; red) cells (2). SOX21 is expressed basal (TRP63; red) and ciliated (FOXJ1; grey) and is high expressed in cells absent of P63 and FOXJ1(→) (3). Scale bar = 25 μm. (C) Schematic overview of human primary bronchial epithelial cell (HPBEC) culture. QPCR analysis of *SOX2* and *SOX21* expression during differentiation of HPBECs on Air-Liquid Interface (ALI). One-way ANOVA (n=4 (ALI cultures of 4 different donors); *** p<0.001). (D) Immunofluorescence analysis of HPBEC 16 days after ALI of SOX2 (red), SOX21 (green), and TP63 (blue) or FOXJ1 (blue). XZ plane shows basal cells (TP63+) at the basal membrane and ciliated cells (FOXJ1+) at the luminal side. Scale bar = 25 μm. The grey bar graphs the MFI of SOX2 or SOX21 in basal and luminal cells. The red bar graphs show the MFI of SOX2 or SOX21 in FOXJ+ and FOXJ-luminal cells. The circled FOXJ1+ nuclei show high expression of SOX2 with sporadic high expression of SOX21(◄) as well. T-test (n=3 (ALI cultures of 3 different donors); *p<0.05, *** p<0.001). (E) Dot-plot of the MFI of SOX2 (y-as: left graph) or SOX21 (y-as: right graph). Each dot represents a cell and ALI cultures of 2 different donors are included. The red filled circles are FOXJ1+ ciliated cells, green filled circles are TP63+ basal cells and non-filled are FOXJ1-TRP63-cells. (F) Schematic overview of in vitro basal cell differentiation. In human, SOX21 levels are high in an intermediate state, SOX21 levels are decreasing when FOXJ1 is present. SOX2 levels are mainly elevated in the end stage of differentiation to ciliated cells (FOXJ1+).

To functionally assess the role of SOX21 in human adult airway epithelium, we isolated airway epithelial cells from human bronchial epithelium. Both SOX2 and SOX21 are expressed throughout the bronchial epithelium (Fig. 6B). ALI cultures showed that, similar to mice, *SOX2* and *SOX21* were also expressed *in vitro*. After prolonged culturing, a significant increase of *SOX2* and a slight increase of *SOX21* was observed (Fig. 6C, Fig. S5B). Both SOX2 and SOX21 were expressed higher in the luminal fraction when compared to basal cells (Fig. 6D). Most basal cells were low in SOX2 and SOX21 expression (88.3%), while only a few basal cells were SOX2^low^ SOX21^high^ (8.3%) and almost no basal cells were SOX2^high^SOX21^low^ or SOX2^high^SOX21^high^ (Fig. S5D). Next, we determined whether the cells on luminal side that were SOX2^high^ and/or SOX21^high^ were FOXJ1+ ciliated cells. Comparing FOXJ1+ and FOXJ1-nuclei showed that ciliated cells were high in SOX2 expression, but SOX21 was highest in non-ciliated cells (Fig. 6D, S5C). Measurement of the fluorescence intensity of SOX2 and SOX21 showed that most ciliated cells were SOX2^high^SOX21^low^ (75%) (Fig. S5D), while the intensities of either SOX2 or SOX21 for ciliated (FOXJ1+), basal (TP63+) and FOXJ1-P63-cells, revealed that SOX21 was mainly high in (a subpopulation of) double negative cells (Fig. 6E). We hypothesize that these cells are intermediate cells transitioning from basal cells to a luminal cell fate, the para-basal cell. In accordance with this observation, sections of human airway epithelium, show SOX21^high^ cells in between the basal and luminal layer, which are cells positive for both basal cell marker KRT5 and luminal cell marker KRT8 (Fig. 6B-2,3, arrows). In addition, we compared our results to published single-cell RNA sequencing data on human primary bronchial epithelial ALI culture (Plasschaert, Zilionis et al. 2018). This confirmed that high levels of *SOX21* mRNA denote an intermediate cell type. Furthermore, similar to our observation, high levels of *SOX2* were observed both in the intermediate state and in differentiated ciliated cells (Fig. S5E). In conclusion, we observe high levels of SOX21 in para-basal cells, which have lost TP63 expression and do not (yet) express FOXJ1. These levels of SOX21 decrease when differentiation continues to ciliated cells. SOX2 levels are mainly elevated in the maturation of differentiated FOXJ1+ ciliated cells (Fig. 6C). In conclusion, our data suggest that SOX21 is an early determinant of differentiation of basal to ciliated cells, while SOX2 is important in the maturation of ciliated cells.

## DISCUSSION

During lung development, a proximal-to-distal epithelial gradient is observed by the separation of proximal SOX2 and distal SOX9 expressing cells. Here we show a further regionalization of the proximal epithelium by marking a SOX2+/SOX21+ proximal zone, zone 1. Within this zone, progenitor cells differentiate to basal cells. With the use of human fetal lung organoids, we confirmed SOX21 expression in SOX2+ progenitor stalk cells (Nikolic, Caritg et al. 2017). In addition, SOX21 becomes more widely expressed when the organoids are differentiated to airway organoids. Based on our findings, we propose that SOX2+SOX21+ expressing progenitor cells in the murine and human fetal lung organoids are progenitor cells in a zone destined for differentiation and that SOX21 is important in balancing the maintenance and differentiation of SOX2+ airway progenitor cells.

SOX2 has been shown to be a key regulator in regulating proliferation and differentiation in many different stem cells populations, however the chromatin regions targeted by SOX2 are cell-type specific. The regulation of stem cells by SOX2 is dependent on its co-factors as well as on its expression levels (Brafman, Moya et al. 2013, Hagey, Klum et al. 2018). We observed a correlation between the high expression of SOX2 and appearance of SOX21 at the dorsal side of the proximal airways and trachea. Additionally, ectopic induction of SOX2 in distal cells resulted in an up-regulation of SOX21. We suggest that SOX2 requires a certain threshold level of expression to induce SOX21. Interestingly, SOX21 expression was first observed at embryonic stage (E10.5-11.5), at a time where cells adopt a proximally restricted fate for the extrapulmonary airways (Yang, Riccio et al. 2018). We suggest that SOX21 is a downstream effector of SOX2, while the extrapulmonary airways are formed, separating them form progenitors of the intrapulmonary airways that are only SOX2 positive. A similar induction of SOX21 by SOX2 has been described in the 4-cell stage embryo, embryonic stem cells (ESCs) and neuronal progenitor cells. The function of SOX21 and its synergy or antagonism of SOX2 activity has been shown to be highly context-dependent (Kopp, Ormsbee et al. 2008, Mallanna, Ormsbee et al. 2010, Chakravarthy, Ormsbee et al. 2011, Goolam, Scialdone et al. 2016). For example, SOX2 maintains the neuronal progenitor state in the developing nervous system and SOX21 can either stimulate differentiation or help maintain the progenitor pool depending on external stimuli and expression levels (Graham, Khudyakov et al. 2003, Ohba, Chiyoda et al. 2004, Sandberg, Kallstrom et al. 2005, Matsuda, Kuwako et al. 2012). Here, we show that SOX9+ distal progenitor cells are maintained upon ectopic expression of SOX21, and differentiation to SOX2+ progenitor cells is not initiated. Thus, SOX21 itself is not capable of driving differentiation to SOX2+ progenitor cells in the presence of distal mesenchymal signaling. However, once coexpressed with SOX2, SOX21 expression associates with the region where SOX2 progenitor cells differentiate to basal cells.

Using *Sox21*^+/-^ and *Sox21*^-/-^ mice, we show that the presence of SOX21 within this zone is important to suppress the differentiation of SOX2+ progenitor cells to basal and ciliated cells. Surprisingly, the number of basal cells is increased upon deletion of SOX21, but also after ectopic expression of SOX21 using a SPC-promotor. This seemingly contradicting result may be caused by the artificial induction of SOX21 in SOX9 expressing cells, which does not induce SOX2. SOX21 may initiate specification of these SOX9+ cells to a proximal phenotype, which become even more determined after these SOX9+ cells exit the influence of the distal mesenchymal FGF10 signaling. The latter causes the cells to express SOX2, leading to a more pronounced induction of the proximal fate. The extension of the SOX2+ SOX21+ zone 1 coincides with an induction of TRP63 basal cells, which might also occur in this SOX21 induced mouse model. The initiation of a proximal cell fate by ectopic expression of SOX21, followed by SOX2 expression may explain why the cysts observed in the SOX21 expressing lungs are smaller than the cysts observed when SOX2, the major proximal cell fate inducer, is expressed.

SOX21 and SOX2 co-expression continues in adult TE and SMGs, both regions where progenitor cells reside. We used naphthalene injury and *in vitro* analysis to study whether SOX21 and SOX2 function are similar in adult progenitors as during development. We show that reduced SOX2 levels inhibit the contribution of SOX9+ SMG cells to TE injury, while reduced SOX21 levels promote their contribution. Furthermore, reduced levels of SOX2 decreased differentiation of basal cells to ciliated cells *in vitro* and *in vivo*, while reduced levels of SOX21 only increased basal cell differentiation *in vitro* but not *in vivo*. The latter might be explained the fact that differentiation of progenitor cells *in vivo* is regulated by an epithelial-mesenchymal interaction, which might inhibit further differentiation to ciliated cells when regeneration is complete (Volckaert, Yuan et al. 2017). Thus, both during development and regeneration of the airway epithelium, SOX21 acts as a suppressor of SOX2+ progenitor cell differentiation, while SOX2 levels are important for stimulating differentiation. Using a luciferase assay, we specifically show that SOX21 can antagonize SOX2 function on certain promotor regions. However, to fully understand the function of SOX21 within SOX2+ airway progenitors, signaling/environmental cues, additional co-factors, and expression levels are likely to play a role.

SOX2 and SOX21 showed a similar but not identical expression pattern in human airway epithelium. As opposed to mouse tracheal epithelium of only a basal and luminal cell layer, the human proximal airway epithelium contains an additional layer of intermediate cells or para-basal cells (Mercer, Russell et al. 1994, Boers, Ambergen et al. 1998). Using human primary bronchial epithelial ALI cultures, we show high levels of SOX21 in TP63-FOXJ1-cells and suggest that these cells represent the intermediate layer of para-basal cells transitioning to a luminal cell fate. Basal cell hyperplasia of the airway epithelium is an important disease feature in smokers, COPD and cystic fibrosis (CF) patients (Rock, Randell et al. 2010). A better understanding on how SOX2 and SOX21 levels control human basal cell proliferation and differentiation may therefore help to identify therapeutic targets of airway remodeling within these patients.

Our data provide a new understanding of proximal-distal patterning of the airways and the regulation of SOX2 progenitor cells within development and regeneration of airway epithelium. De-regulation of SOX2 and SOX21 expression levels can alter branching morphogenesis and differentiation of airway epithelium. We show that SOX21 is a suppressor of differentiation when SOX2 expression levels are high and when progenitor cells are prone to differentiate.

## MATERIALS AND METHODS

### Mice

All animal experimental protocols were approved by the animal welfare committee of the veterinary authorities of the Erasmus Medical Center. Mice were kept under standard conditions. Mouse strains *SPC-rtTA* (gift of Jeffrey Whitsett), *pTT::myc-SOX2*, *Sox2-CreERT* (Jackson Labs, stock number 017593) and *SOX21-KO* (gift of Stavros Malas) have been described (Gontan, de Munck et al. 2008, Kiso, Tanaka et al. 2009, Arnold, Sarkar et al. 2011). A murine *Sox21 cDNA* with a N-terminal myc epitope was subcloned in a modified pTRE-Tight (Clontech) vector (pTT::myc-sox21). Pronuclear microinjection of a linearized fragment was performed to develop transgenic mice, and three independent lines were tested. To induce expression of *Sox2* or *Sox21* during lung development, the *pTT::myc-Sox2* or *pTT::mycSox2*1 mice were crossed with SPC-rtTA mice and doxycycline was administered to dams in the drinking water (2 mg/ml doxycycline, 5% sucrose) from gestational day 6.5 onwards. Wild-type animals were C57BL/6.

### Naphthalene Injury

Adult mice (∼8-12 weeks) were injured with a single intraperitoneal injection of 300 mg/kg naphthalene. Naphthalene (Sigma; 184500) was freshly prepared and dissolved in corn oil (Sgima; C8267). Corn oil injection served as baseline control. Groups of mice were sacrificed 2, 5 and 20 days post injury (DPI) (number of mice per group is indicated in each figure).

### Mouse tracheal epithelial cell culture

Mouse tracheal epithelial cell (MTEC) culture was performed as previously described (Eenjes, Mertens et al. 2018). Briefly, MTECs were isolated from mice adult trachea and cultured in KSFM expansion medium (Table 1) on collagen coated plastic (50 μg/cm^2^ of rat tail collagen Type IV (SERVA, 47256.01) in 0.02N acetic acid(Sigma; 537020)). After expansion, 8*10^4^ MTECs were plated per collagen coated 12-well insert (Corning Inc, Corning, USA) in proliferation medium (Table 1) for air-liquid interface (ALI) culture. When confluent, MTECs were exposed to air by removing proliferation medium and adding MTEC differentiation medium (Table 1) to the lower chamber. MTECs were cultured in standard conditions; at 37°C in a humidified incubator with 5% CO_2_.

**TABLE 1:** Medium

| KSFM-hPBEC medium |  |  |  |
| --- | --- | --- | --- |
| Reagent | Company | Cat. No. | Final Concentration |
| KSFM | Gibco | 17005034 | n/a |
| Penicillin / Streptomycin | Lonza | DE17-602e | 100 U/ml 100 µg/ml |
| Bovine Pituitary Extract | Gibco | 13028014 | 0.03 mg/ml |
| Human EGF | Peprtech | 315-09 | 25 ng/ml |
| Isoproterenol | Sigma | I-6504 | 1 µM |
| KSFM-MTEC Expansion medium |  |  |  |
| Reagent | Company | Cat. No. | Final Concentration |
| KSFM | Gibco | 17005034 | n/a |
| Penicillin / Streptomycin | Lonza | DE17-602e | 100 U/ml 100 µg/ml |
| Bovine Pituitary Extract | Gibco | 13028014 | 0.03 mg/ml |
| Mouse EGF | Peprtech | 315-09 | 25 ng/ml |
| Isoproterenol | Sigma | I-6504 | 1 µM |
| Rock Inhibitor (Y27632) | Axon MedChem | 1683 | 10 µM |
| DAPT | Axon MedChem | 1484 | 5 µM |
| MTEC Proliferation Medium |  |  |  |
| Reagent | Company | Cat. No. | Final Concentration |
| DMEM:F12 | Gibco | 1133032 | n/a |
| Penicillin / Streptomycin | Lonza | DE17-602e | 100 U/ml 100 µg/ml |
| NaHCO<sub>3</sub> | Gibco | 25080094 | 0.03% (w/v) |
| Fetal Calf Serum | HyClone | SH30071.03 | 5% |
| L-Glutamine | Gibco | 25030081 | 1.5 mM |
| Insulin-Transferin-Selenium | Gibco | 41400045 | 1x |
| Cholera Toxin | Sigma | C8052 | 0.1 µg/ml |
| Bovine Pituitary Extract | Gibco | 13028014 | 0.03 mg/ml |
| Mouse EGF | Peprtech | 315-09 | 25 ng/ml |
| Rock Inhibitor (Y27632) | Axon MedChem | 1683 | 10 µM |
| Retinoic Acid | Sigma | R2625 | 0.05 µM |
| MTEC Differentiation Medium |  |  |  |
| Reagent | Company | Cat. No. | Final Concentration |
| DMEM:F12 | Gibco | 1133032 | n/a |
| Penicillin / Streptomycin | Lonza | DE17-602e | 100 U/ml 100 µg/ml |
| NaHCO<sub>3</sub> | Gibco | 25080094 | 0.03% (w/v) |
| Bovine Serum Albumin | Gibco | 15260037 | 0.1% (w/v) |
| L-Glutamine | Gibco | 25030081 | 1.5 mM |
| Insulin-Transferin-Selenium | Gibco | 41400045 | 1x |
| Cholera Toxin | Sigma | C8052 | 0.025 µg/ml |
| Bovine Pituitary Extract | Gibco | 13028014 | 0.03 mg/ml |
| Mouse EGF | Peprotech | 315-09 | 5 ng/ml |
| Retinoic Acid | Sigma | R2625 | 0.05 μM |
| **Human Foetal Organoid medium** |  |  |  |
| **Reagent** | **Company** | **Cat. No.** | **Final Concentration** |
| Advanced DMEM:F12 | Invitrogen | 12634-034 | n/a |
| R-Spondin | Peprotech | 120-38 | 500 ng/ml |
| Noggin | Peprotech | 120-10C | 100 ng/ml |
| Fgf10 | Peprotech | 100-26 | 100 ng/ml |
| Fgf7 | Peprotech | 100-19 | 100 ng/ml |
| EGF | Peprotech | AF-100-15 | 50 ng/ml |
| CHIR 99021 | Stem Cell Techn. | 72052 | 3 μM |
| SB 431542 | Tocris | 1614 | 10 μM |
| B27 supplement (- VitA) | ThermoFisher | 12587-010 | 1x |
| N-Acetylcysteine | Sigma | A9165 | 1.25 mM |
| Glutamax 100x | Invitrogen | 12634-034 | 1x |
| N2 | ThermoFisher | 17502-048 | 1x |
| Hepes | Gibco | 15630-56 | 10 mM |
| Penicillin / Streptomycin | Lonza | DE17-602e | 100 U/ml 100 μg/ml |
| Primocin | Invivogen | Ant-pm-1 | 50 μg/ml |
| **Human Adult Organoid medium** |  |  |  |
| **Reagent** | **Company** | **Cat. No.** | **Final Concentration** |
| Advanced DMEM:F12 | Invitrogen | 12634-034 | n/a |
| R-Spondin | Peprotech | 120-38 | 500 ng/ml |
| Noggin | Peprotech | 120-10C | 100 ng/ml |
| Fgf10 | Peprotech | 100-26 | 100 ng/ml |
| Fgf7 | Peprotech | 100-19 | 25 ng/ml |
| SB202190 | Sigma | S7067 | 500 nM |
| A83-01 | Tocris | 2939 | 500 nM |
| Y-27632 | Axon MedChem | 1683 | 5 μM |
| B27 supplement | Gibco | 17504-44 | 1x |
| N-Acetylcysteine | Sigma | A9165 | 1.25 mM |
| Nicotinamide | Sigma | N0636 | 5 mM |
| Glutamax 100x | Invitrogen | 12634-034 | 1x |
| Hepes | Gibco | 15630-56 | 10 mM |
| Penicillin / Streptomycin | Lonza | DE17-602e | 100 U/ml 100 μg/ml |
| Primocin | Invivogen | Ant-pm-1 | 50 μg/ml |

### Human primary epithelial cell culture

Human primary airway epithelial cell (HPBEC) culture was performed as previously described (Amatngalim, Schrumpf et al. 2018). Lung tissue was obtained from residual, tumor-free, material obtained at lung resection surgery for lung cancer. The Medical Ethical Committee of the Erasmus MC Rotterdam granted permission for this study (METC 2012-512).

Briefly, cells were isolated from healthy bronchial tissue by incubation in 0.15% Protease XIV (Sigma; P5147) for 2hrs at 37 °C. The inside of the bronchi was scraped in cold PBS (Sigma; D8537) and the obtained airway cells were centrifuged and resuspended in KSFM-HPBEC medium for expansion (table 1). Culture plates were coated with 10 μg/mL human fibronectin (Millipore; FC010), 30 μg/mL BSA (Roche; 1073508600) and 10 μg/mL PureCol (Advanced Biomatrix; 5005-B) for 2 hrs at 37°C. Upon confluency, HPBECs were frozen (4*10^5^ cells / vial) and stored for later use.

When used for ALI culture, cells were thawed and seeded in a coated 10cm-dish, grown until confluent in KSFM-HPBEC medium, trypsinized and 8*10^4^ of HPBECs were plated per 12-transwell insert (Corning Inc, Corning, USA). On inserts, the HPBECs were cultured in bronchial epithelial growth medium (ScienCell Research Laboratories, Carlsbad, USA; 3211). Basal bronchial epithelial growth medium was first diluted 1:1 with DMEM (Gibco; 41966). Next, 1x supplement and 1x pen/strep (Lonza; DE17-602e) was added to 500 mL (BEGM). Retinoic acid (1 nM, RA) was freshly added. When cells reached full confluency, the BEGM medium was removed from the apical chamber and only supplied to the basal chamber and freshly supplemented with 50 nM RA. The medium was changed every other day and the apical chamber was rinsed with PBS. HPBECs were cultured in standard conditions; at 37°C in a humidified incubator with 5% CO_2_.

### Human fetal organoid culture

The culture of human fetal lung (17 weeks of gestation) and adult airway organoids was performed as previously described (Nikolic, Caritg et al. 2017, Miller, Hill et al. 2018).

Lung lobes were dissociated using dispase (Corning; 354235) on ice for 30 minutes. Tissue pieces were then incubated in 100% FBS on ice for 15 minutes, after which they were transferred to a 10% FBS solution in Advanced DMEM with Glutamax, P/S and Hepes. Lung bud tips were separated from mesenchymal cells through repeated pipetting. Pieces of tissue were resuspended in 30 μl of basal membrane extract (BME type 2, Trevigon; 3533-010-02), transferred to a 48-well plate and incubated at 37°C to solidify the BME. After 5 min, 300 μl of self-renewing fetal lung organoid medium (table 1) was added (Nikolic, Caritg et al. 2017). Medium was refreshed every 3-4 days. Every 2 weeks, organoids were split 1:3 to 1:6. The medium was aspirated and cold PBS was added to the well to re-solidify the BME. Organoids were disrupted using a 1000 μl tip with on top a 2 μl tip. The disrupted organoids were centrifuged at 300 g for 5 min, resuspended in BME and re-plated. To differentiate to airway epithelium, fetal lung organoids were split, resuspended in BME and replated. To initiate differentiation of fetal lung organoids to airway organoids, human adult organoids medium was added after splitting. Organoids were grown under standard culture conditions (37 °C, 5% CO_2_).

### Immunofluorescence

#### Tissue

Mouse embryonic lungs and human bronchial tissue were fixed overnight in 4% PFA (Sigma; 441244) at 4 °C. Post-fixation, samples were washed with PBS, de-hydrated to 100% ethanol, transferred to xylene and processed to paraffin wax for embedding.

Organoids were retrieved from the BME by adding cold PBS to the 48 well. Organoids were centrifuged at 150g for 5 min and fixed overnight in 4% PFA at 4 °C. Post-fixation, organoids sink to the bottom of the tube. Centrifuging was avoided after fixation to keep the organoids intact. The organoids were embedded in 4% low-melting agarose. The organoids were washed in PBS for 30 min and manually de-hydrated by 50 min incubation steps in 50%, 70%, 85%, 95% and 2 times 100% ethanol. Organoids were further processed by 3 times xylene for 20-30 min, washed 3 times for 20-30 min in 60 °C warm paraffin to remove all traces of xylene. The organoids were placed in a mold and embedded in paraffin. Paraffin blocks were sectioned at 5 μm and dried overnight at 37 °C.

Sections were deparaffinized by 3 times 3 min xylene washes, followed by rehydration in distilled water. Antigen retrieval was performed by boiling the slides in Tris-EDTA (10M Tris, 1M EDTA) buffer pH=9.0 for 15 min. Slides were cooled down for 30 min and transferred to PBS. For SOX21 staining, the Tyramide Signal Amplification (TSA) kit was used (Invitrogen, B40922, according to manufacturer’s protocol). When using the TSA kit, a hydrogen peroxide (35%) blocking step was performed after boiling. Sections were blocked for 1 hr at room temperature (RT) in 3% BSA (Roche; 10735086001), 0.05% Tween (Sigma; P1379) in PBS. Primary antibodies (table 2) were diluted in blocking buffer and incubated with the sections overnight at 4 °C. The next day, sections were washed 3 times for 5 min at room temperature (RT) in PBS with 0.05% Tween. Secondary antibodies (table 2) were added in blocking buffer and incubated for 2 hrs at RT. DAPI (4’,6-Diamidino-2-Phenylindole) solution (BD Pharmingen, 564907, 1:4000) was added to the secondary antibodies for nuclear staining. After incubation, 3 times 5 min washes in PBS-0.05% Tween and one wash in PBS was performed, sections were mounted using Mowiol reagent (For 100 mL: 2,4% m/v Mowiol (Sigma; 81381), 4,75% m/v glycerol, 12 % v/v Tris 0.2M pH=8.5 in dH_2_O till 100 mL). All sections were imaged on a Leica SP5 confocal microscope.

**TABLE 2:** Antibodies

| Primary antibodies |  |  |  |  |
| --- | --- | --- | --- | --- |
| Antibody | Species | Company | Cat. No. | Dilution |
| β-TUBULIN IV | Mouse | BioGenex | MU178-UC | 1:100 |
| FOXJ1 | Mouse | eBioscience | 14-9965 | 1:200 |
| KRT5 | Rabbit | Biolegend | Poly19055 | 1:500 |
| KRT8 | Rat | DSHB | TROMA-I | 1:100 |
| MYC (Immunofluorescence) | Rabbit | Abcam | AB9106 | 1:500 |
| SCGB1A1(CCSP, uteroglobin) | Rabbit | Abcam | AB40873 | 1:200 |
| SCGB3A1 / HIN-1 | Mouse | R&D systems | AF1790 | 1:200 |
| SCGB3A2 / UGRP1 | Goat | R&D systems | AF3465 | 1:500 |
| SMOOTH MUSCLE ACTIN | Mouse | Neomarkers | MS-113-P1 | 1:500 |
| SOX2 | Rabbit | Seven-Hills | WRAB-SOX2 | 1:500 |
| SOX2 | Goat | Immune systems | GT15098 | 1:500 |
| SOX21 | Goat | R&D systems | AF3538 | 1:100 (TSA: 9 min) |
| SOX9 | Rabbit | Abcam | AB185230 | 1:500 |
| TRP63 | Mouse | Abcam | AB735 | 1:100 |
| TRP73 | Rabbit | Abcam | AB40658 | 1:100 |
| FLAG | Rabbit | Sigma | F7425 | 1:1000 |
| MYC (Western Blot) | Rabbit | Abcam | AB9106 | 1:1000 |
| β -TUBULIN | Mouse | Sigma | T8328 | 1:2000 |

| Secondary antibodies |  |  |  |
| --- | --- | --- | --- |
| Antibody | Company | Cat. No. | Dilution |
| Alexa Fluor ® 405, 488, 594<br>Donkey anti Goat IgG | Jackson ImmunoResearch | 705-475-147,<br>705-545-147,<br>705-585-147 | 1:500 |
| Alexa Fluor ®488, 594, 647<br>Donkey anti Mouse IgG | Jackson ImmunoResearch | 715-545-151,<br>715-585-151,<br>711-605-151 | 1:500 |
| Alexa Fluor ®488, 594, 647<br>Donkey anti Rabbit IgG | Jackson ImmunoResearch | 711-545-152,<br>711-585-151,<br>711-605-152 | 1:500 |
| Alexa Fluor ®488, 594<br>Donkey anti Rat IgG | Jackson ImmunoResearch | 712-545-150<br>712-585-153 | 1:500 |
| Anti-Goat HRP conjugated | Jackson ImmunoResearch | 705-035-147 | 1:500 |

#### Air-liquid interface culture

Human or mouse ALI cultures were washed with PBS and fixed on inserts in 4% PFA at RT for 15 min. Inserts were then washed 3 times for 5 min in 0.3% TritonX (Simga; T8787) in PBS and blocked for 1 hr at RT in 5% normal donkey serum (NDS, Millipore; S30), 1% BSA 0.3% TritonX in PBS. Primary antibodies (table 2) were diluted in blocking buffer and incubated overnight at 4 °C. The next day, inserts were 3 times rinsed with 0.03% TritonX in PBS followed by 3 washes for 10 min at RT in PBS with 0.03% TritonX. Secondary antibodies (table 2) were added in blocking buffer and incubated for 2 hrs at RT. DAPI solution (BD Pharmingen, 564907, 1:2000) was added to the secondary antibodies as well. After incubation, inserts were 3 times rinsed with 0.03% TritonX in PBS followed by 3 washes for 10 min at RT. Inserts were covered by a coverslip using Miowol reagent. Images were collated on a Leica SP5 confocal microscope.

### Image analysis

#### Fluorescence intensity measurements

Intensity measurements in MTEC and HPBEC cultures were performed on 3 separate isolations from wild-type mice or donors and measured using ImageJ. Of each n, more that 500 nuclei were manually selected on the DAPI staining. In each nucleus, the intensity of SOX2 and SOX21 was measured. The MFI for each n and each intensity measurement was calculated by dividing it by the average intensity of that measurement in the same n.

#### Counting

To standardized counting between animals, basal and ciliated cells were counted during lung development in a square of 400 μm around the first branch at the medial side of the bronchi. In this way, we could determine a position in the *SOX21*-/- animal where in the wildtype SOX21 is highest expressed. Of each genotype and each n, 3 sections were counted and the percentage of ciliated and basal cells were calculated based on the total number of airway epithelial (SOX2+) cells.

Five days’ after Naphthalene injury, the number of basal (TRP63+), ciliated (FOXJ1+), dividing (KI67+) and SOX9+ cells were counted in the tracheal epithelium from cartilage ring (C) 0 till C1. Twenty days’ post injury, basal, ciliated and dividing cells were counted from C0 till C6. Of each animal 3 sections were counted throughout the trachea.

Of the MTEC culture, the number of FOXJ1+ nuclei were counted per 775 μm^2^. For determining, the differentiation to cilia and secretory cells, the percentage of TUBIV+ and SCGB3A1+ area per 775 μm^2^ was measured. The number of TRP63+ basal and TRP73+ cells were counted respectively to the number of nuclei present in each field. Each n are separate isolations of different animal, and per n, 5 fields of 775 μm^2^ were counted.

### RNA isolation, cDNA synthesis and qRT-PCR analysis

Human or mouse airway cells were removed from the insert by scraping them off the insert into cold PBS. Cells were collected in an Eppendorf tube and centrifuged at 800g for 5 min at 4 °C. PBS was aspirated and the cell pellet was snap frozen in liquid nitrogen and stored at −80 °C till RNA isolation.

To isolate RNA, 500 μl of TRI Reagent^®^ (Sigma, T9424) was added to the cell pellet. RNA extraction was performed according to the TRI Reagent ^®^ protocol. RNA concentrations were measured using NanoDrop (ThermoFisher Scientific). First strand cDNA synthesis was synthesized using 2 μg RNA, MLV Reverse transcriptase (Sigma, M1302) and Oligo-dT primers (self-designed: 23xT+A, 23xT+C and 23xT+G). For one qRT-PCR reaction, 0.5 μl of cDNA was used with Platinum Taq polymerase (Invitrogen, 18038042) and SybrGreen (Sigma, S9430). The primer combinations for the qRT-PCR are listed in table 3. Normalized gene expression was calculated using the ddCT method relative to GAPDH (mouse) or B-ACTIN (human) control.

**TABLE 3:**
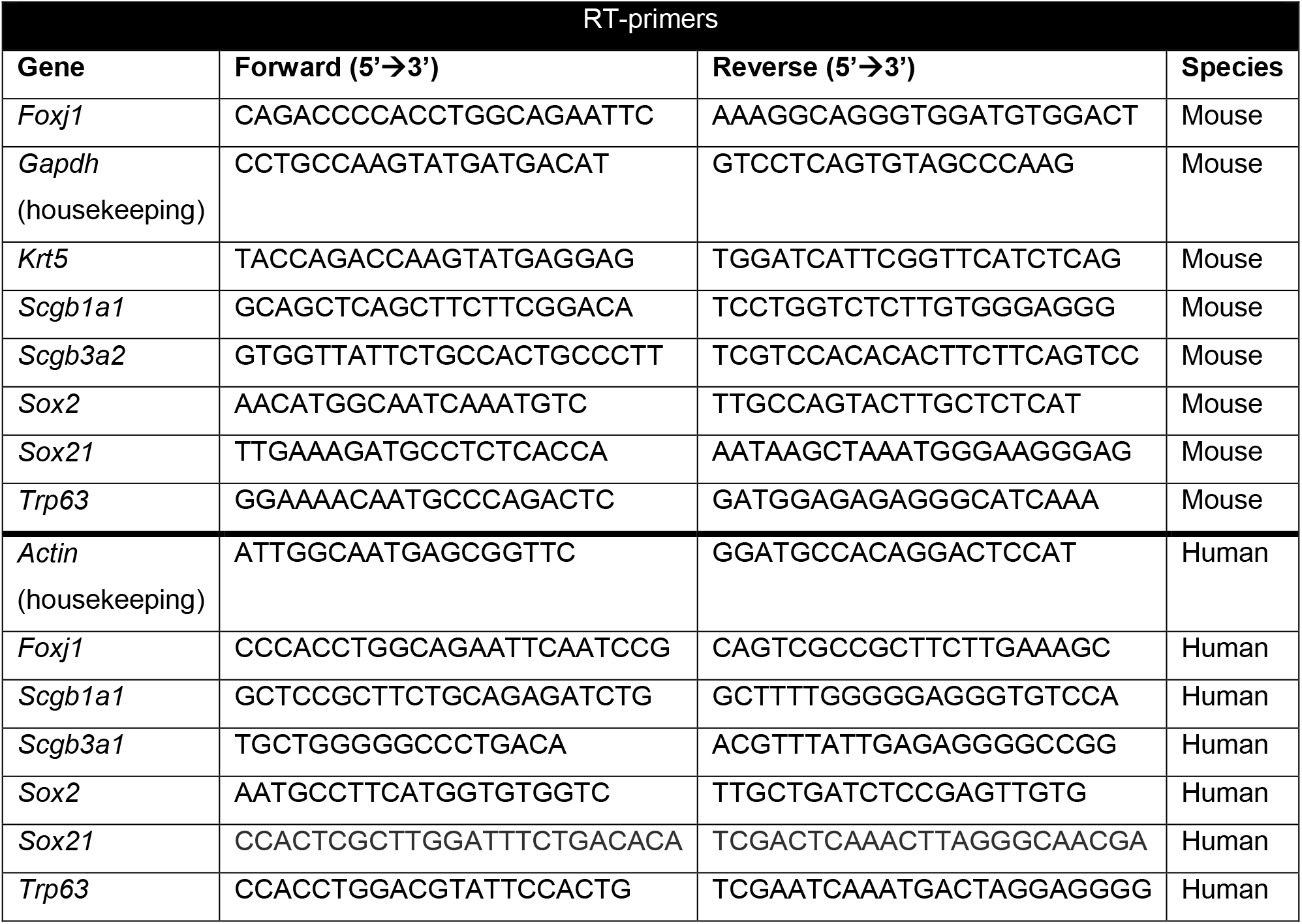
RT-PCR primers

### Luciferase Assay

#### Cloning

Promotor regions were PCR-amplified (primers listed in table 4) from mouse genomic DNA. To each primer, a restriction site was added, to clone the promotor region into the pGL4.10 [luc2] construct (Promega; E6651). The promotor sequence included the sequence of a transcriptional active area, which was identified with a RNA polymerase II ChIP in mouse tracheal epithelial cells (Marshall, Mays et al. 2016). We used *in silico* analysis to predict SOX21 binding sites (MatInsepctor, GenomatixSuite v3.10 and http://jaspar2016.genereg.net/).

**TABLE 4:**
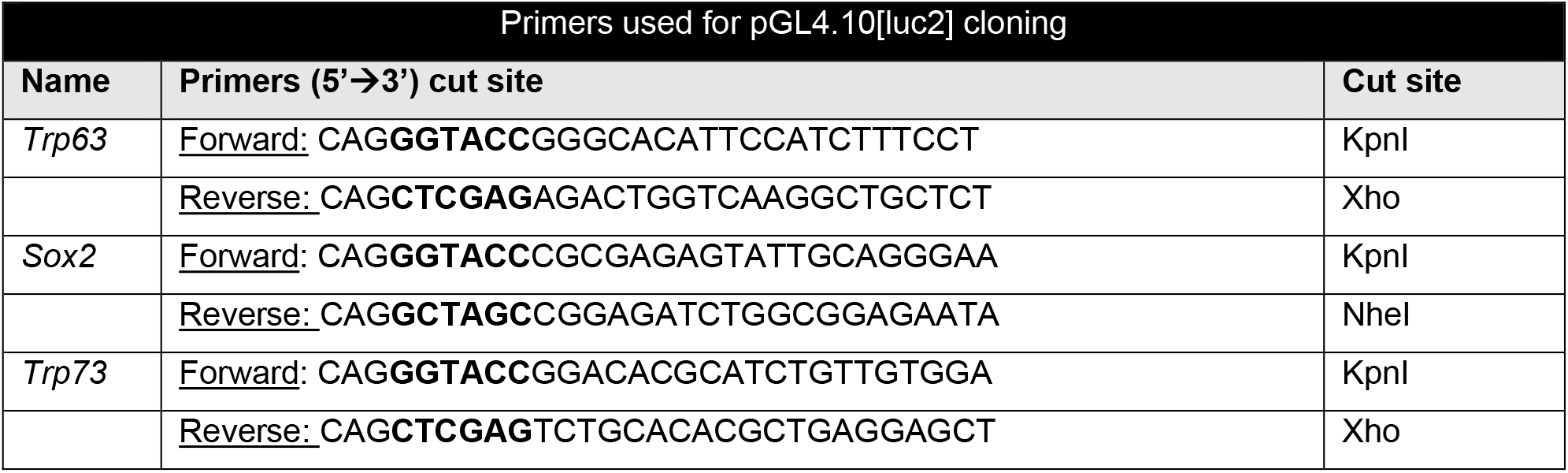
Primers used for pGL4.10[luc2] cloning

#### Luciferase Assay

HELA cells were plated in a 12-well plate in DMEM+ 10%FCS and transiently transfected using Lipofectamine3000 (Thermofisher, L3000001). Each transfection consisted of 250 ng expression plasmid (pcDNA3-control (Addgene; n/a), pcDNA3-Sox2FLAG (homemad), pcDNA3-Sox21MYC (homemad)), 250 ng pGL4.10[luc2] reporter plasmid and 2.5 ng TK-Renilla plasmid (Promega; E2241) (transfection control). Luciferase activity was measured 48 hrs after transfection using the Dual-Luciferase^®^ Reporter Assay System (Promega, E1910). Plate reader VICTOR™ X4 was used to measure Firefly and Renilla luminescence. Firefly luminescence of each sample was calculated by dividing the Firefly luminiscence by the Renilla luminescence. The increase or decrease of luciferase activity was then normalized to the pGL4.10[luc2] reporter plasmid transfected with the pcDNA3-control expression plasmid.

#### Western blot

Samples were prepared from cell lysates used for luciferase measurements. Cells were lysed in the lysis buffer that was included in the Dual-Luciferase^®^ Reporter Assay System (Promega, E1910). To the cell lysate, 8M urea (to denature the DNA), 50mM 1,4-dithiothreitol (DTT, Sigma) and 1x SDS Sample buffer was added. Samples were boiled and loaded on a 12% SDS-polyacrylamide gel and blotted onto a PVDF membrane (Immobilon^®^-P transfer membrane, Millipore). The blots were blocked for 1 h in PBS containing 0.05% Tween-20 and 3% BSA at room temperature, and probed overnight with primary antibodies at 4 °C (Table 2). Next day, membranes were washed three times with PBS containing 0.05% Tween-20 and incubated for 1 h with horseradish peroxidase (HRP)-conjugated secondary antibodies (DAKO) at a dilution of 1:10,000. Signal was detected with Amersham™ ECL™ Prime Western Blotting Detection Reagent (GE Healthcare). Blots were developed using the Amersham Imager 600GE (GE Healthcare).

### Statistics

Statistical analysis was performed using Prism5 (Graphpad). For all measurement, three or more biological replicates were used. Data are represented as means ± standard error of mean (SEM) with the data points present in each graph. Statistical differences between samples were assed with Student unpaired t-test, one-way ANOVA (post-test: Tukey) or two-way ANOVA (post-test: Bonferroni). P-values below 0.05 are considered significant. The number of replicates and statistical tests used are indicated in the figure legends.

### Single Cell RNA sequencing of HPBEC

Expression levels of *SOX2* and *SOX21* in ALI cultures of HPBEC were analyzed using the dataset previously published (Plasschaert, Zilionis et al. 2018). The data set was available on the following link: https://kleintools.hms.harvard.edu/tools/springViewer_1_6_dev.html?datasets/reference_HB_ECs/reference_HBECs

## ACKNOWLEDGEMENT

We thank, Mart Lamers and Bart Haagmans (Department of Viroscience, Erasmus MC) for assistance and supplying the human fetal lung organoids, Thomas Koudstaal (department of Pulmonary Medicine, Erasmus MC) for supplying lung resection material for the isolation of human primary bronchial epithelial cells, and Frank Grosveld for critically reading the manuscript (Department of Cell Biology, Erasmus MC). This work was supported by a grant from the Sophia Foundation for Medical Research (grant number S14-12).

## AUTHOR CONTRIBUTIONS

E.E. designed, performed, analyzed the experiments and wrote the paper. M.V.B.K, A.D.M and L.B.K performed the experiments and reviewed the manuscript. J.M.S reviewed the manuscript. D.T. funding acquisition and reviewed the paper. J.J.P. wrote, reviewed and edited the paper. R.J.R. supervision, funding acquisition, experimental design and wrote the paper.

---

**Sup. Figure 1:**
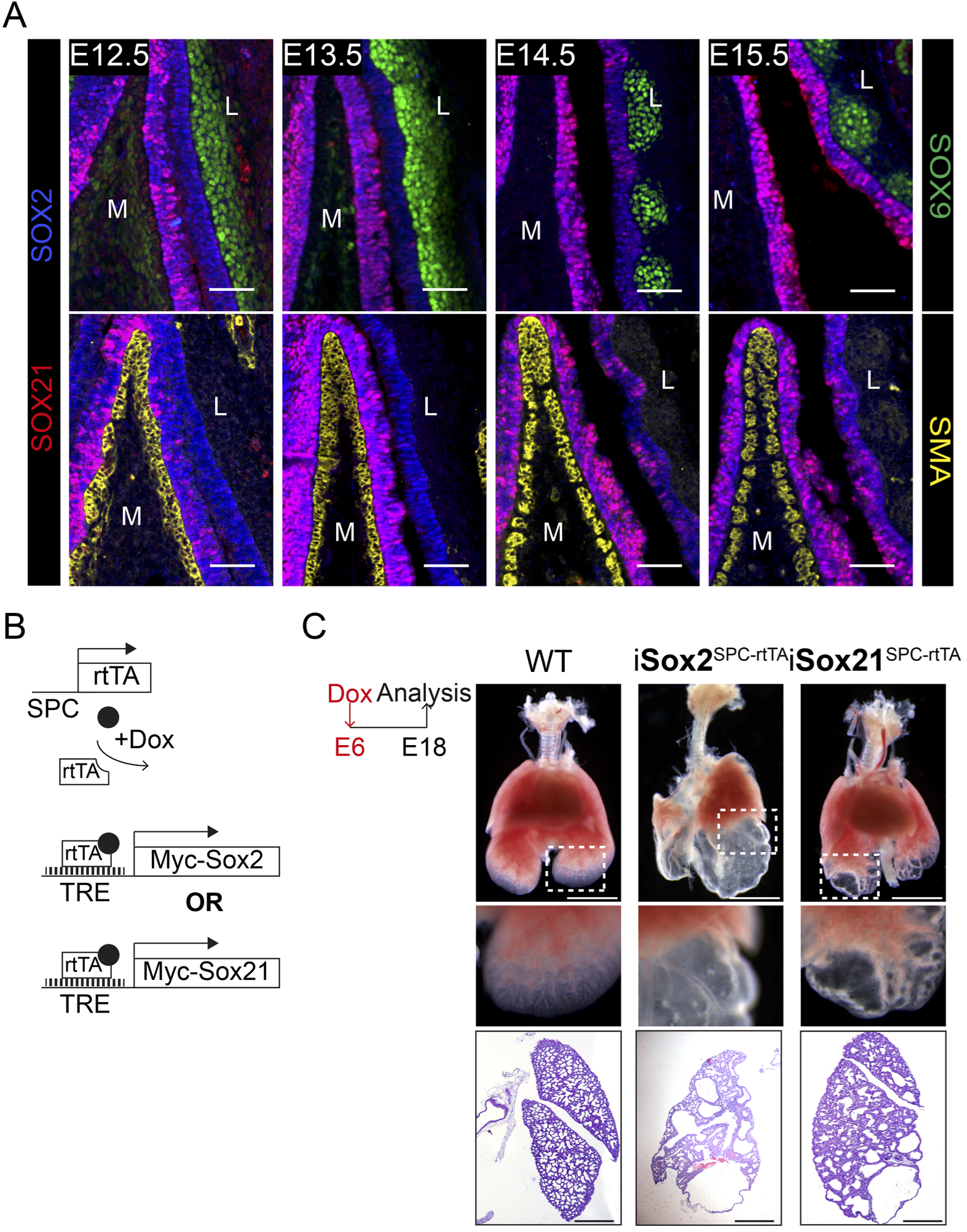
SOX21 is unilateral detected in the developing trachea and overexpression of SOX21 leads to lung cysts. (A) Immunostaining on sections of the main bronchi immediately distal of the carina from mice at embryonic day (E) 12.5, 13.5, 14.5, and 15.5 using SOX2 (blue), SOX21 (red) and either SOX9 (green, top row) or SMA (yellow, bottom row). The mesenchymal cells on the medial (M) side express Smooth Muscle Actin (SMA, yellow). At the lateral (L) side, mesenchymal cells express SOX9 (green). Scale bar = 50 μm. (B) Schematic representation of the iSox2^SPC-rtTA^ and iSox21^SPC-rtTA^ mouse models. (C) Bright field images of E18.5 lungs from control, iSox2^SPC-rtTA^ and iSox21^SPC-rtTA^ mice that received doxycycline from E6 onwards. Scale bar = 2 mm. HE staining of the lungs show the size of the cysts that are present in the lungs of iSox2^SPC-rtTA^ and iSox21^SPC-rtTA^ mice. Scale bar = 0.5 mm.

**Sup. Figure 2:**
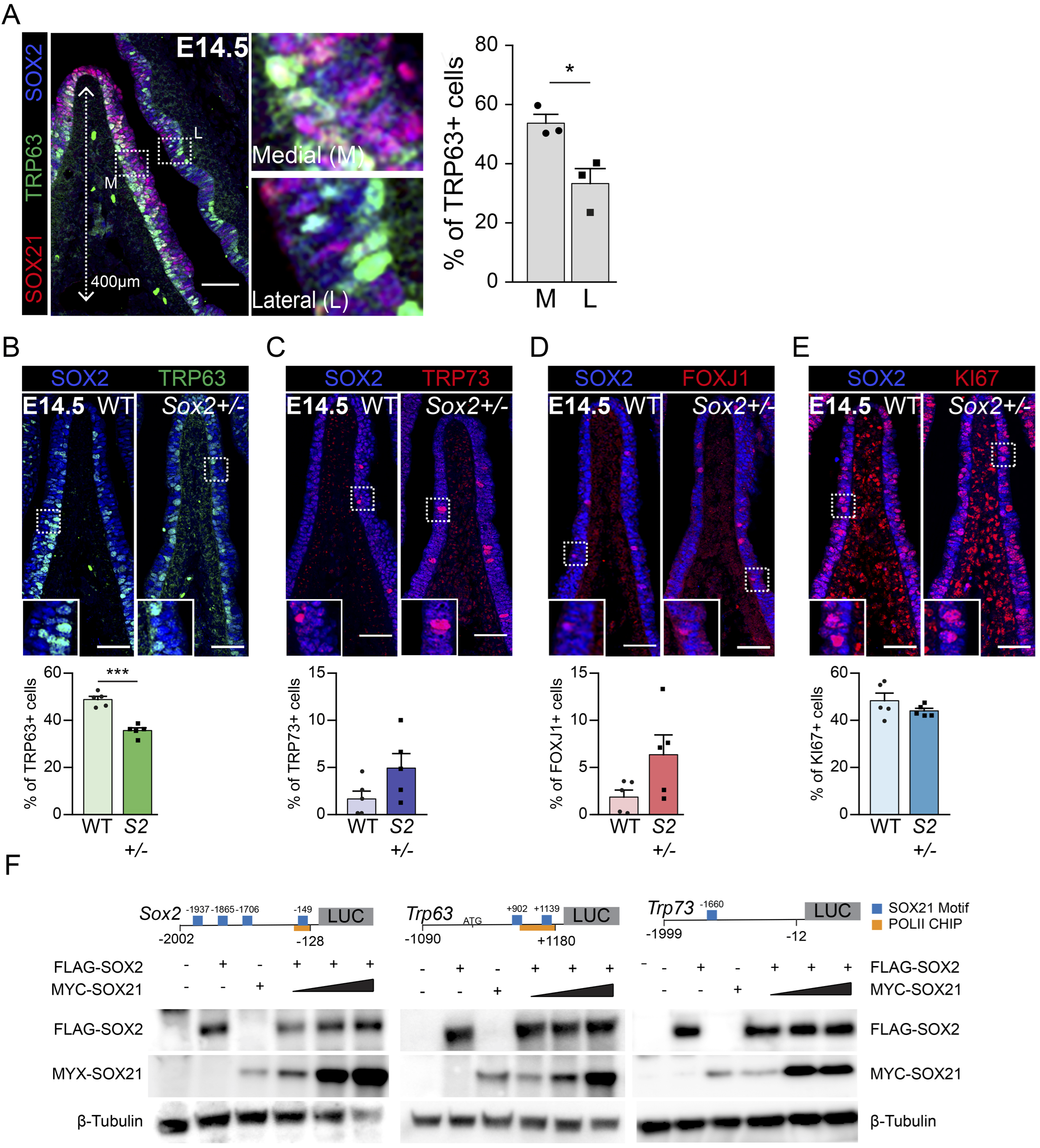
Basal cells arise in the SOX2+ SOX21+ region. (A) Co-staining of TRP63 (green) for basal cells, SOX2 (blue) and SOX21 (red) on longitudinal sections of the main bronchi at E14.5. The graph shows the percentage of basal cells in medial versus lateral side of the bronchi 400 μm proximal of the first branch. T-test (n=3; *p<0.05, *** p<0.001). M = medial, L = lateral. Scale bar = 100 μm. (B-E) Immunofluorescence and quantification of the number of TRP63+ basal cells (B), TRP73+ cells (C), FOXJ1+ ciliated cells (D) and KI67+ dividing cells (E) at E14.5 in wildtype (WT) and Sox2+/- mice. The number of cells were counted within the first 400 μm immediately distal of the carina at the medial side of the airway. T-test (n=5; * p<0.05, *** p<0.001). Scale bar = 50 μm. (G) Western blots showing the protein levels of transfected FLAG-SOX2 and MYC-SOX21 of the luciferase assay belonging to figure 3E. Beta-tubulin was used as loading control.

**Sup. Figure 3:**
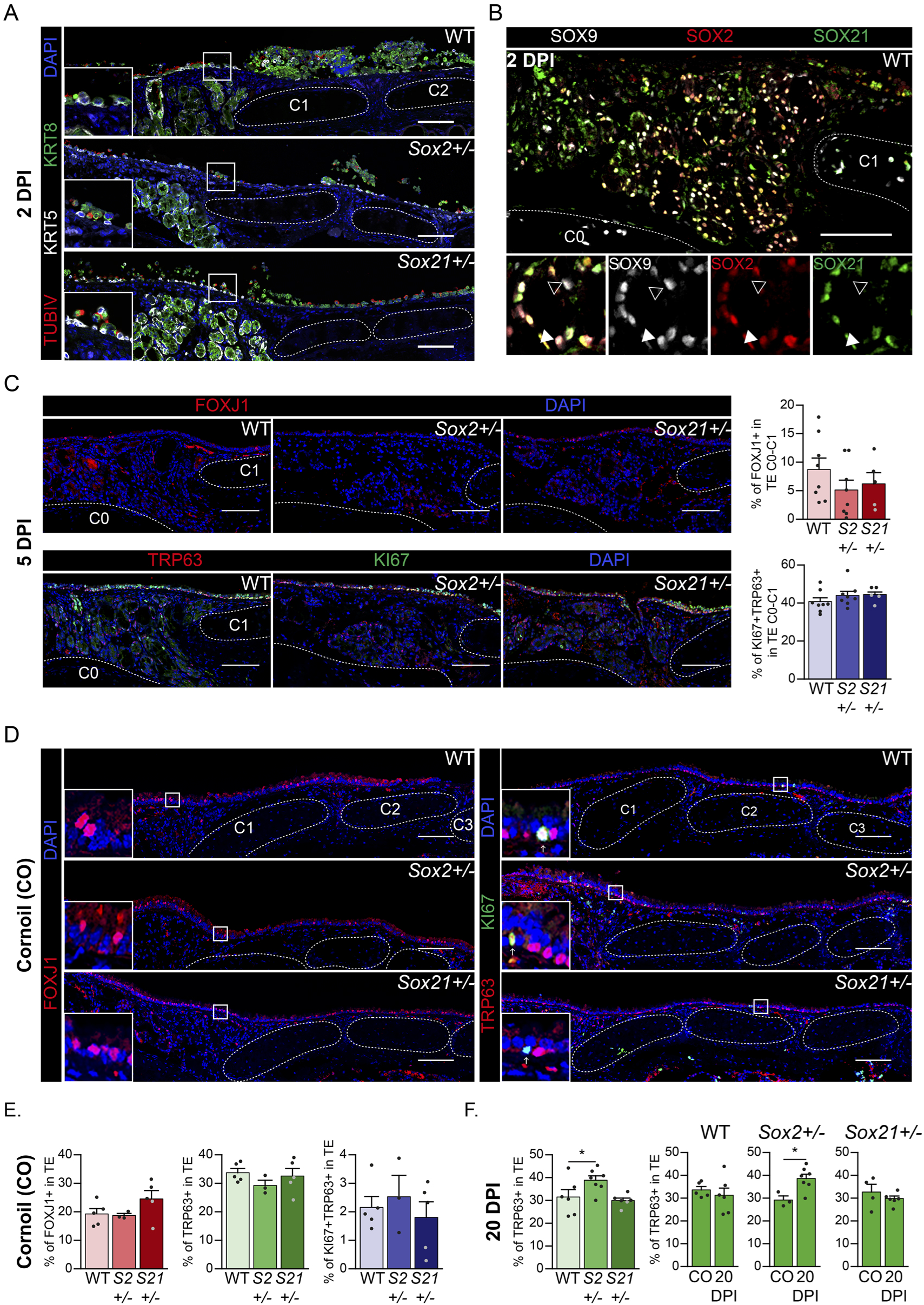
Regeneration is delayed in SOX2 deficient tracheal epithelium. (A) Immunofluorescence staining on tracheal sections of wildtype (WT), Sox2+/- and Sox21+/-, 2 days post injury (2 DPI) of Keratin 5 (KRT, grey), Keratin 8 (KRT8, green), TubilinIV (T∪BIV, red). (B) Tracheal section showing Submucosal Glands (SMGs) at 2 DPI of wildtype mice. SOX2 (red) and SOX21 (green) expression is detected in SOX9+ SMG cells. Closed white arrowheads indicate cells only expressing SOX2 and SOX9, open arrowheads indicate cells only expressing SOX9. Scale bar = 100 μm. Scale bar = 100 μm. C = Cartilage ring. (C) Immunofluorescence of FOXJ1+ ciliated cells (top row) orTRP63÷ KI67+ dividing basal cells (bottom row), 5 DPI in wildtype (WT), Sox2+/- and Sox21+/- mice. Scale bar = 100 μm. Quantification of the number of FOXJ1+ ciliated cells from CO till C1 between genotypes. One-way ANOVA (WT n = 5, Sox2+/- n = 3, Sox21+/- n = 5, * p<0.05) (D) Immunofluorescence of FOXJ1+ ciliated cells (left) orTRP63÷ KI67+ dividing basal cells (right), on tracheal sections of cornoil exposed (CO) mice. Scale bar = 100 μm. (E) Quantification of the number of FOXJ1+ ciliated cells, basal cells or dividing basal cells from CO till C6 between genotypes in cornoil exposed (CO) mice. One-way ANOVA (WT n = 8, Sox2+/- n= 8, Sox21+/- n = 5) (F) Quantification of the number of basal cells from C0 till C6 between genotypes. One-way ANOVA (WT n = 6, Sox2+/- n = 7, Sox21+/- n = 6, * p<0.05). Quantification of basal cells, 20 DPI compared to CO animals of each WT, Sox2÷/- and Sox21+/-. T-test (WT: CO n = 5 and 2ODPI n = 6, Sox2÷/-: CO n = 3 and 20DPI n = 7, Sox21+/-: CO n = 4 and 20DPI n = 6, * p<0.05).

**Sup. Figure 4:**
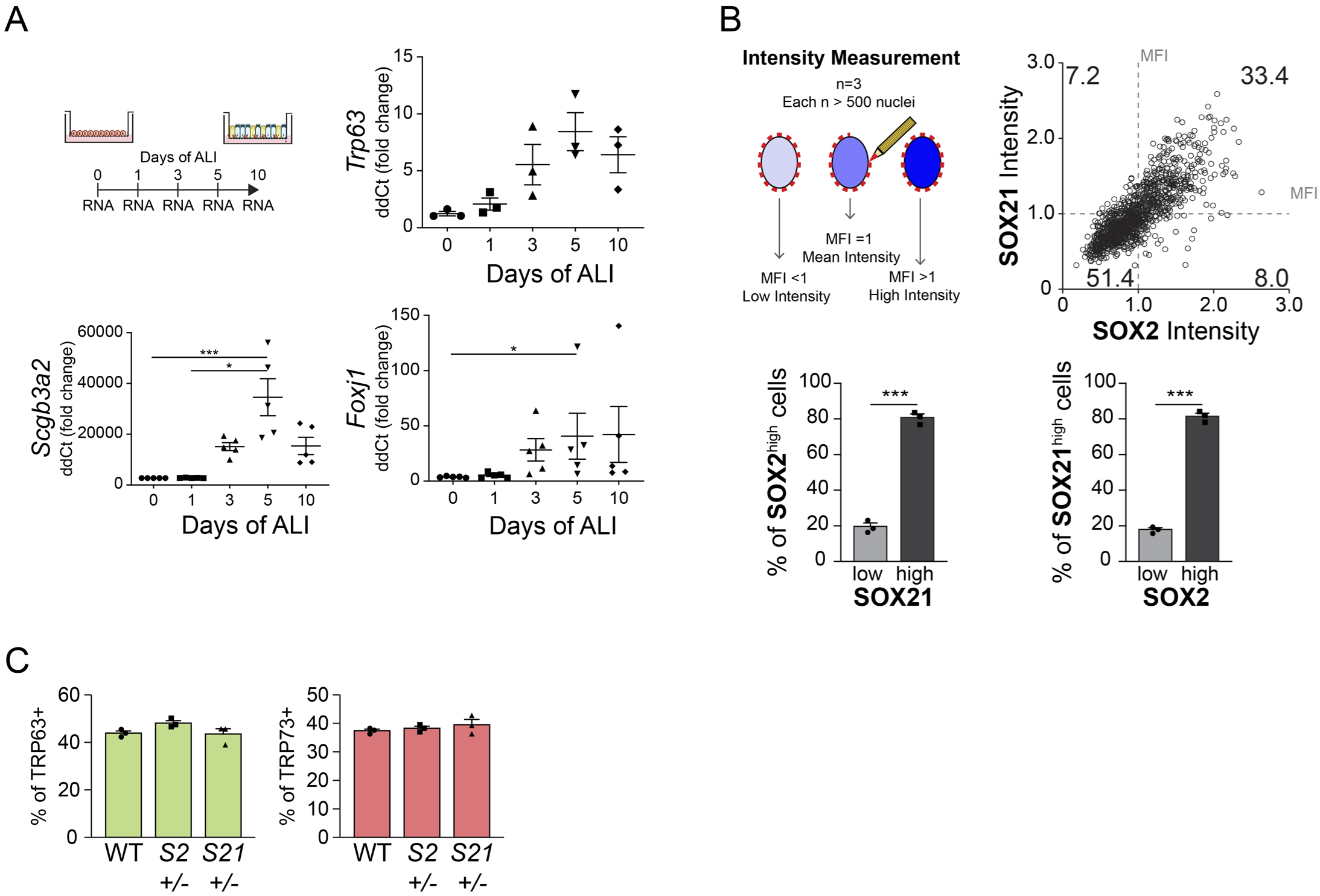
SOX2 and SOX21 are inversely correlated with basal cell differentiation to ciliated cells. (A) Schematic overview of mouse tracheal epithelial cell (MTEC) culture experiment. QPCR analysis of TRP63, SCGB3A2 and FOXJ1 expression during differentiation of MTECs on Air-Liquid Interface (ALI). One-way ANOVA (n=5; * p<0.05, *** p<0.001). (B) Schematic representation of the method to measure the mean fluorescence intensity (MFI) on day 10 of ALI. Dot-plot of the MFI of SOX2 (x-as) and SOX21 (y-as). The number represents the percentage of cells in each quadrant of the total. Bar graph is the quantification of SOX2 high expressing cells (MFl> 1) that are either high (MFI>1) or low (MFI<1) in expression of SOX21 (left), and SOX21 high expressing cells (MFI>1) that are either high (MFI> 1) or low (MFl< 1) in expression of SOX2 (right). T-test (n=3; *** p<0.001). (C) Quantification of the percentage of all TRP63+ basal cells and all TRP73+ cells belonging to figure 4E. One-way ANOVA (n=3).

**Sup. Figure 5:**
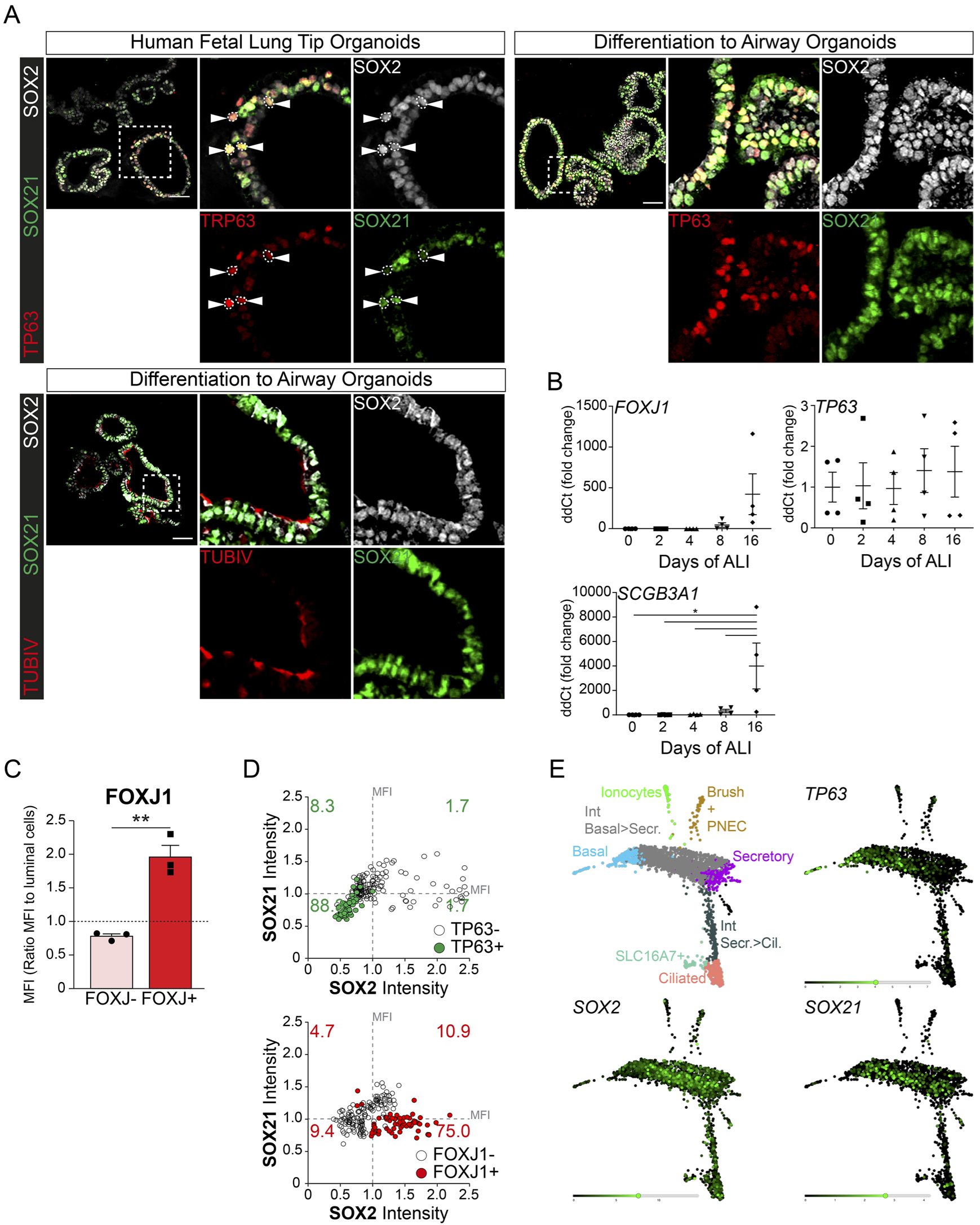
SOX21 is expressed in human airway epithelium. (A) Immunofluorescence analysis of SOX2 (grey), TRP63 (red) and SOX21 (green) in fetal lung organoids. Closed arrowheads (►) show cells positive for SOX2, TRP63 and SOX21. Fetal lung organoids differentiated to airway epithelium as shown by the presence of ciliated cells (TUBIV; red) or basal cells (TP63, red) have an abundant expression of SOX21 (green). Scale bar = 50 μm. (B) QPCR analysis of TP63, SCGB3A1 and FOXJ1 expression during differentiation of HPEBCs on Air-Liquid Interface (ALI). One-way ANOVA(n=4; *** p<0.001). (C) Bar graphs shows the separation of luminal cells in FOXJ1+ versus FOXJ1-cells, belonging to figure 6D. A higher MFI was found in the cells selected for ciliated cells. T-test (n=3 (ALI cultures of 3 different donors); **p<0.005). (D) Dot-plot representation of the MFI of SOX2 (x-as) and SOX21 (y-as). The green filled circles are TP63+ basal cells (upper graph) and the number in each quadrant represents the percentage of basal cells in each square. The red filled circles are FOXJ1+ ciliated cells (lower graph) and the number in each quadrant represents the percentage of ciliated cells in each square. Each dot represents a cell and ALI cultures of 2 different donors are included. (E) Previous published single cell RNA sequencing data on HPBEC ALI culture showing the expression levels of TRP63, SOX2 and SOX21 in the different cell populations (Plasschaert, Zilionis et al. 2018).

